# Feature Selectivity for Circadian Regulation within Melanopsin Photoreceptors

**DOI:** 10.64898/2025.12.30.695276

**Authors:** Elliott S. Milner, Hannah A. Blume, Navid Mousavi, Michael C. Brown, Franklin Caval-Holme, Nguyen-Minh Viet, Takeharu Nagai, Michael Tri H. Do

## Abstract

The day-night cycle drives profound environmental changes, which animals anticipate using the circadian clock. The clock synchronizes with this cycle by extracting features of light that indicate solar elevation. It uses a specialized dynamic range to sense relevant photon rates and precise, extended temporal integration to encode total photon counts. This feature selectivity is distinct from that of visual perception and is thought to arise in the brain. We have found it within the retina’s M1 melanopsin photoreceptors, which signal light to the clock. M1s have heterogeneous intrinsic and synaptic drives, and collectively account for the dynamic range of circadian photoregulation. Both individually and as a population, M1s approach the level of temporal integration observed behaviorally. M1s and behavior match closely when the pupillary light reflex, another M1-driven function, dynamically shapes retinal illumination. Thus, the earliest steps of photoreception are tuned to signifiers of Earth’s rotation.

---

The day-night cycle drives variations in critical parameters that include visibility, the level of ultraviolet radiation, temperature, and the types of active species. Organisms have evolved a circadian clock to anticipate these variations^1^. As the day-night cycle changes, the clock maintains alignment by sensing light. It selects for features of illumination that reflect the Sun’s position in the sky. For example, the clock has a dynamic range that matches relevant intensities^2,3^, with a threshold that is far higher than that of visual perception^4^. This allows visually-guided behavior without clock resetting. The clock also responds to the total number of photons counted over many minutes^5,6^. This smooths away fluctuations that are unrelated to time of day, such as the transient darkening from a passing cloud. By contrast, visual perception resolves millisecond timing differences. There is evidence that the clock’s hallmark dynamic range and temporal integration arise within the brain, including in the principal circadian clock— the suprachiasmatic nucleus (SCN)—and its inhibitory inputs^7,8^.

The clock senses light through intrinsically photosensitive retinal ganglion cells (ipRGCs)^9^. These ocular neurons capture photons directly, using the melanopsin photopigment, and receive signals originating with the rod and cone photoreceptors. IpRGCs are essential for light’s regulation of the clock, and those of the M1 type are sufficient^9,10^.

M1s have heterogeneous, intrinsic tunings to light intensity, with melanopsin-driven dynamic ranges that differ by orders of magnitude in the intensities covered^11–13^. Whether M1 synaptic inputs covary is unknown, even though they are necessary for the clock’s normal sensitivity to light^2^. Indeed, the field lacks consensus on the photosensitivities of M1 intrinsic responses and extrinsic drives^11,13–15^. Furthermore, the kinetics of rod, cone, and melanopsin responses differ over orders of magnitude, from tens of milliseconds to minutes^13,16,17^. How such responses collectively support the clock’s temporal integration is unknown. Here, we ask how rod, cone, and melanopsin signals combine to drive M1 responses, and how those responses align with the clock’s distinct feature selectivity.

## RESULTS

### Genetic identification of cells with minimal photoexcitation

Behavioral experiments indicate that rods, signaling through ipRGCs, set the threshold of circadian photoregulation^2^. Rods are desensitized by conventional methods of identifying ipRGCs and other cell types, which use fluorescence^18^. We sought an alternative in bioluminescence, testing nano-lantern (NL)^19^. NL uses a substrate rather than excitation light as energy for signaling. We engineered a Cre-dependent NL mouse line (**Extended Data Fig. 1**) and crossed it with one expressing Cre from the melanopsin gene locus (*Opn4^Cre^*)^20^. Cells of varied morphology were bioluminescent, consistent with recognized ipRGC types (M1-M6; **Extended Data Fig. 2**). We examined M4s because they are among the most sensitive RGCs and bioluminesce in our conditions, thus providing a conservative test case^21^. M4 firing rates were transiently elevated following substrate exposure. Nevertheless, responses to flashes and steps of light showed no detectable desensitization at any time point (**Extended Data Fig. 2** and **Extended Data Table 1**). Thus, even the most photosensitive RGCs can be identified quite innocuously with NL.

### M1 ipRGCs at threshold are driven by rod signals

To enrich for M1 ipRGCs that directly influence the clock, we injected NL-encoding viruses into the hypothalamus, centering on the SCN (**Fig. 1a**)^13,22^. Labeled M1s fired spontaneously in darkness at a low rate (2.8 ± 3.3 Hz, 32 cells; **Extended Data Fig. 2g**). Flashes modulated firing at an intensity of 0.50 ± 0.73 log R*/rod, ∼200-fold higher than that needed for M4s (**Extended Data Fig. 2f** and **Extended Data Table 1**). Modulation was brief for most M1s, indicating drive by synaptic input^13^. Thus, at threshold, M1s appear to be driven by retinal circuits of moderate sensitivity.

**Fig. 1.**
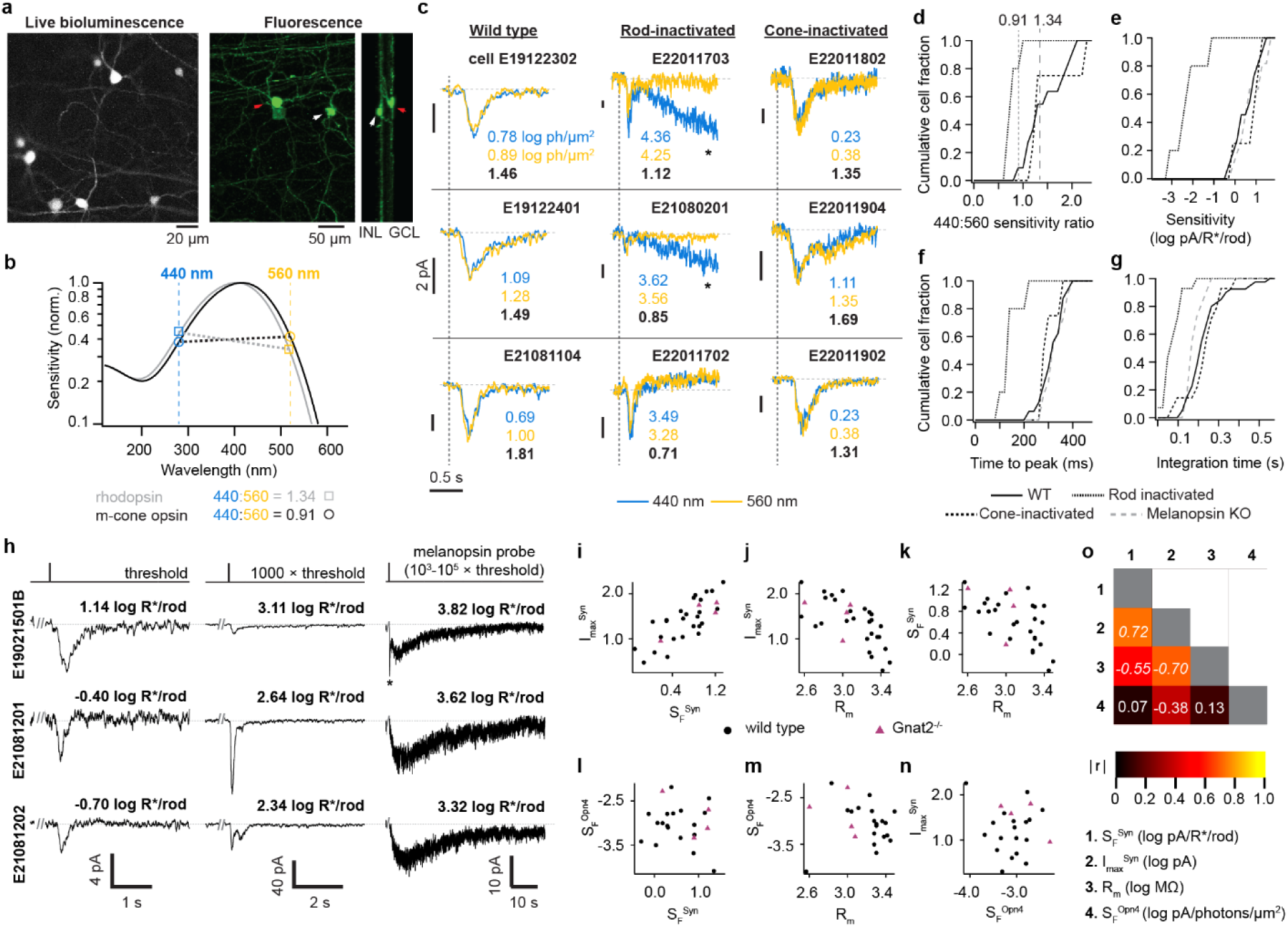
Synaptic and intrinsic drives of M1 ipRGCs. **a.** IpRGCs retrograde-transduced with nano-lantern from the suprachiasmatic nucleus. *Left,* live, bioluminescence visualization. *Right,* Projections of a confocal volume through fixed tissue, showing two cells with M1 morphology: one whose soma is in the Ganglion Cell Layer (red arrowhead) and one in the Inner Nuclear Layer (white arrowhead). **b.** Spectral sensitivities of rhodopsin (gray) and medium-wavelength-sensitive (M) cone photopigment (black). Their sensitivities to 440- and 560-nm light (dashed vertical lines) have ratios of 1.34 and 0.91, respectively. **c.** Example, matched-amplitude responses to 440-nm (blue) and 560-nm (gold) flashes of wild- type (left column), *Gnat1^−/−^* (center), and *Gnat2^−/−^* (right) M1s. Horizontal dashed lines mark 0 pA; vertical dashed lines mark the onset of a 30-ms flash of 440 or 560 nm. Noted are the flash intensities (log photons/μm^2^) and 440:560 sensitivity ratio (black). In *Gnat1^−/^*^−^ retinas, high-intensity 440-nm flashes sometimes evoked melanopsin responses (asterisks), which returned to baseline within ∼60-70 s. Traces are averages of >10 sweeps (500-Hz low-pass filtered for display). -80 mV, 23 °C for stability. **d-g**. Cumulative distributions of 440:560 sensitivity ratio (**d**), absolute flash sensitivity (**e**), time to peak (**f**), and integration time (**g**; normalized integral of the response)^16,17^ across genotypes (11, 5, 4, and 11 cells from wild-type, *Gnat1^−/−^*, *Gnat2^−/−^*, and *Opn4^Cre/Cre^* animals; no *Opn4^Cre/Cre^* in **d**). Dashed and dotted vertical lines in **d** mark the expected spectral sensitivity ratio for rhodopsin and M cone photopigment, respectively. **h**. Sample responses from three M1s (rows) to a flash near threshold (30 ms; left column, average of >10 sweeps); 1000× brighter (30 ms; center column, average of 1-3 sweeps); and even brighter to produce a melanopsin dim-flash response (30-200 ms; right column, single sweep). -80 mV, 500-Hz low-pass filtered for display. Flash intensities are given (R*/rod). **i-n**. Plots of four response parameters (S_F_^Syn^, synaptic flash threshold sensitivity; I ^Syn^, synaptic response amplitude at 1000× threshold; R_m_, membrane resistance; and S_F_^Opn4^, melanopsin dim-flash sensitivity) measured in single M1s. **o**. Correlation coefficients for the four parameters (data from wild type only). Spearman r values that were statistically significant are italicized (p ≤ 0.0167 with Bonferroni correction for multiple comparisons).

To identify the origin of these threshold responses, we tested spectral sensitivity (**Fig. 1b**). We measured currents evoked by flashes of two wavelengths, adjusting their intensities to obtain just-detectable responses of similar amplitudes (**Fig. 1c**). Dividing each threshold response amplitude by flash intensity gives sensitivity. The ratio of sensitivities between wavelengths indicates rod drive (**Fig. 1d**). Rod drive is also indicated by the high sensitivity and slow kinetics of responses. With rods inactivated, spectral sensitivity, sensitivity, and kinetics are consistent with cone drive. These parameters appear unchanged with cone inactivation or melanopsin gene deletion (**Fig. 1e-g** and **Extended Data Table 2**). Therefore, the most sensitive photocurrents in M1s originate with rods; we observed no responses consistent with cone or melanopsin drive in the scotopic range.

### Combining synaptic and intrinsic drives within M1 ipRGCs

We delivered flashes to evoke threshold synaptic current (**Fig. 1h** and **Extended Data Fig. 3**). We then delivered more intense flashes, starting at 1000× threshold and going as high as 4.95 log R*/rod to evoke threshold melanopsin current. From these currents, we obtained the sensitivity of synaptic (S_F_^Syn^) and melanopsin (S_F_^Opn4^) drives. We found that S_F_^Syn^ was higher than S_F_^Opn4^ by >4000-fold (mean for 26 cells). Furthermore, S_F_^Syn^ covaried with synaptic currents evoked at 1000× threshold and with input resistance, while S_F_^Opn4^ did not (**Fig. 1i-o**). Thus, synaptic drive appears more orderly than melanopsin drive. However, synaptic drive is not uniform across cells. Flashes far above threshold (1000×) evoked measurable responses ranging from ∼2 to ∼200 pA (**Extended Data Fig. 3**). These observations—that synaptic input is highly variable and uncorrelated with melanopsin sensitivity on a cell-by-cell basis—raise the question of how synaptic and intrinsic drives combine to shape M1 spike outputs.

### Intensity encoding by M1 ipRGCs with synaptic and intrinsic drives

We exposed individual M1s to stepwise illumination of increasing intensity while examining their spike outputs, first under physiological conditions where both synaptic and intrinsic drives are intact, then with synaptic antagonists to isolate the latter (**Fig. 2a**). We confirmed the stationarity of intrinsic drive across repeated stimuli (**Extended Data Fig. 4**). In physiological conditions, the mean threshold intensity that modulated steady firing (I_θ_) by ≥1 Hz was 1.47 ± 0.60 log R*/rod/s (39 cells, **Extended Data Table 3**). In synaptic antagonists, I_θ_ trended higher, to 1.83 ± 0.94 log R*/rod/s (p = 0.054, 30 cells with measurable I_θ_, out of 32). I_θ_ values spanned a range that was ∼8.5-fold broader in antagonists: ∼4,900- versus ∼575-fold. The range in antagonists is a lower bound because 2 cells did not modulate their steady firing even at the highest intensity tested, so their I_θ_ was undefined. Synaptic input appears to regularize the thresholds of M1s, potentially allowing the population to signal with higher fidelity when light is scarce.

**Fig. 2.**
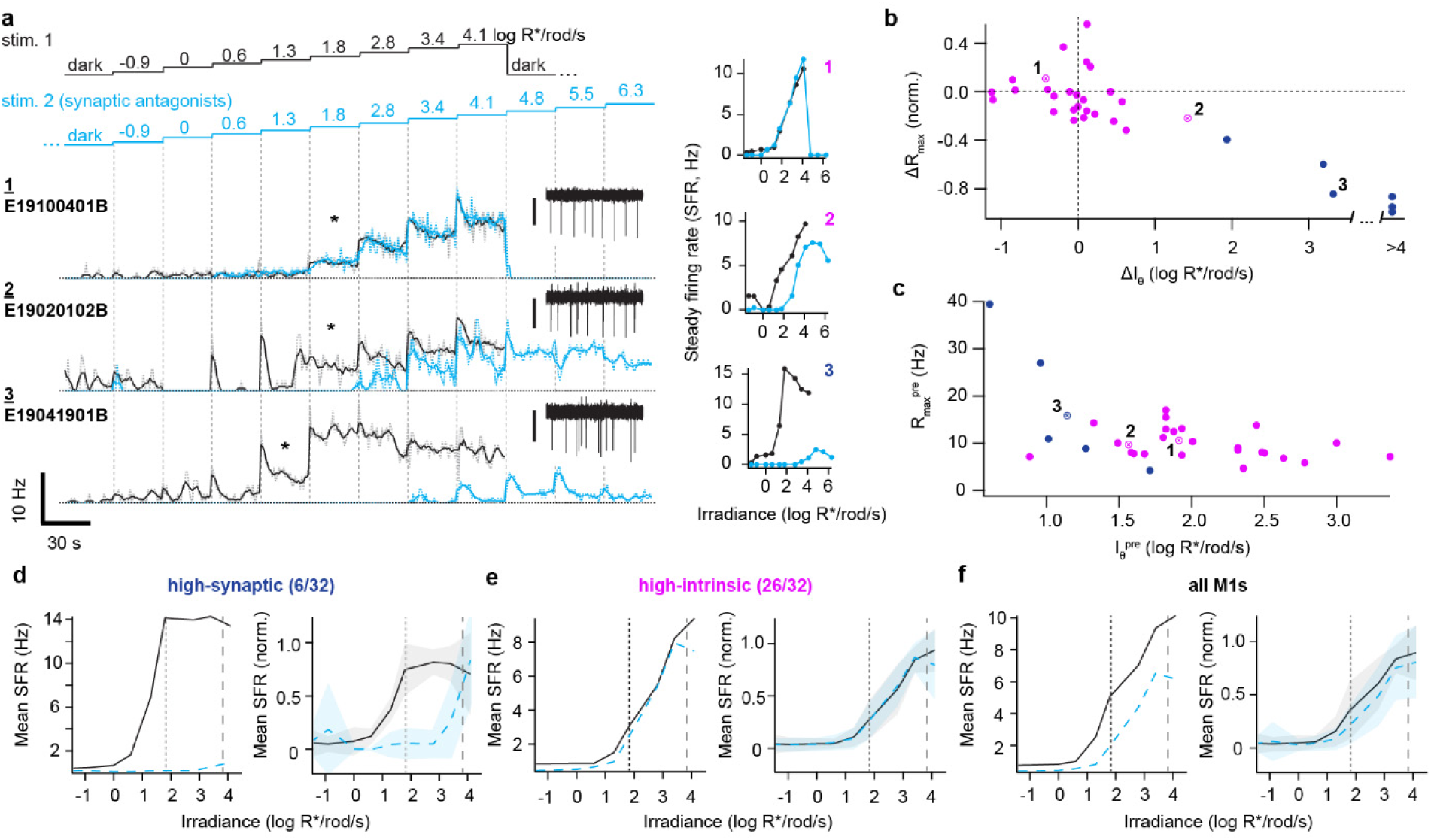
Diverse synaptic and intrinsic drives across M1 ipRGCs account for circadian photoentrainment at and above threshold. **a.** *Left*, Spike rates of three example M1s (rows) during a staircase stimulus (460 nm). 1-s bins and a 5-s moving average are represented by light/dotted lines and dark/solid lines, respectively. Insets show 3 s of raw recordings taken during one step (asterisks) of each cell’s first staircase presentation; scale bars: 40 pA (**1**) and 30 pA (**2,3**). *Right*, Intensity-firing (I-F) relations for the three example cells. Loose-patch recordings, 35 °C. **b.** Difference in maximum steady firing rate (ΔR_max_, normalized to the pre-antagonist firing rate) versus the difference in intensity threshold (ΔI_θ_) between physiological and synaptic antagonist conditions. Clustering gave a “high-synaptic” group (blue; 6 of 32 cells) and a “high-intrinsic” group (magenta; 26 of 32). Sample cells from **a** are marked; cell 3 is high-synaptic, whereas cells 1 and 2 are high-intrinsic. **c.** Maximum steady firing rate plotted against threshold intensity for M1s preceding the addition of synaptic antagonists (R_max_ vs. I_θ_; same cells as in **b**). **d.** *Left*, Average I-F relations before (solid black) and after (dashed cyan) addition of synaptic antagonists for high-synaptic M1s (6 cells). *Right*, Normalized I-F relations (0 minimum to 1 maximum for each cell). Shaded regions are ±1 SD. The intensities at which >80% of wild-type and *Gnat1^−/−^* mice photoentrained^2^ are marked by vertical dotted and dashed lines, respectively. **e**-**f**. As in **d**, but for high-intrinsic M1s (**e**, 26 cells) and all M1s (**f**).

Comparing physiological and antagonist conditions, differences in I_θ_ were correlated with differences in the maximum steady firing rate (R_max_) across all M1s examined (Spearman r = - 0.68, p = 2×10^−4^), suggesting that synaptic input influences M1s across their dynamic range. Nevertheless, ΔI_θ_ and ΔR_max_ were small for most M1s and drastic for a minority. Plotted by these parameters, cells formed a continuum with two densities (**Fig. 2b**). To examine this variation further, we considered cells in terms of heuristic, “high-intrinsic” and “high-synaptic” groups (using k-means clustering; **Methods**). In physiological conditions, high-synaptic cells had lower I_θ_, as expected from stronger rod drive, and tended to have higher R_max_ (**Fig. 2c, Extended Data Table 3**). Variation in the degree of rod drive is also evident in melanopsin-null M1s: Steady firing was practically absent in some (suggesting a normally high dependence on intrinsic photosensitivity) and tracked intensity across orders of magnitude in others (revealing a high degree of sustained synaptic input; **Extended Data Fig. 5**). These data indicate that some M1s act more like autonomous photoreceptors and others more like conventional RGCs at the level of their spike outputs, even though all receive rod-driven synaptic currents at absolute threshold (**Fig. 1**).

### M1 ipRGCs account for the dynamic range of circadian photoentrainment

The threshold of circadian photoentrainment is estimated to be ∼0.37 log R*/rod/s and relies on rod signals carried by ipRGCs^2,3^. High-synaptic M1s are activated at this intensity, raising their steady firing rate (**Extended Data Table 3**). This activity depends on synaptic input, as the firing rate modulation is undetectable in antagonists (**Fig. 2d**). Furthermore, high-intrinsic M1s do not appear to modulate their firing at this intensity even in physiological conditions (**Fig. 2e, Extended Data Table 3**). Therefore, the M1 activity that sets photoentrainment threshold appears to be concentrated in a small subset of the population rather than broadly distributed.

At an intensity of 1.86 log R*/rod/s, most mice photoentrain^2^, likely due to input from both high-synaptic and high-intrinsic M1s (**Fig. 2d-f, Extended Data Table 3**). At even higher intensities, high-synaptic M1s provide little additional signal because their firing rates saturate; meanwhile, high-intrinsic M1s continue to increase firing (**Fig. 2d-e**). Hence, synaptic drive of the most sensitive M1s can account for behavioral threshold, and the progressive recruitment of additional M1s and intrinsic photosensitivity can explain how photoentrainment strengthens as light intensifies.

With rods inactivated, 100-fold higher intensity (∼3.86 log R*/rod/s) is required to photoentrain most mice^2^. In synaptic antagonists at this intensity, high-intrinsic M1s fire strongly while high-synaptic cells fire weakly (**Extended Data Table 3**). Therefore, M1 heterogeneity can explain long-standing behavioral data on photoentrainment thresholds of different genotypes^2^, with the critical synaptic and intrinsic drives being differentially distributed across the population.

### Temporal integration by M1s approaches that of circadian phase shifting

Nelson and Takahashi discovered that animals shift their circadian phase by counting photons precisely over an extended timescale^5^ (**Fig. 3**). We adopted their stimulus paradigm to ask how M1s contribute to this temporal integration. We delivered a given photon count over different durations (30 ms, 3 s, 30 s, and 300 s) while measuring spiking. To estimate the synaptic input to each cell, we also measured threshold flash responses (**Extended Data Fig. 2f**).

**Fig. 3.**
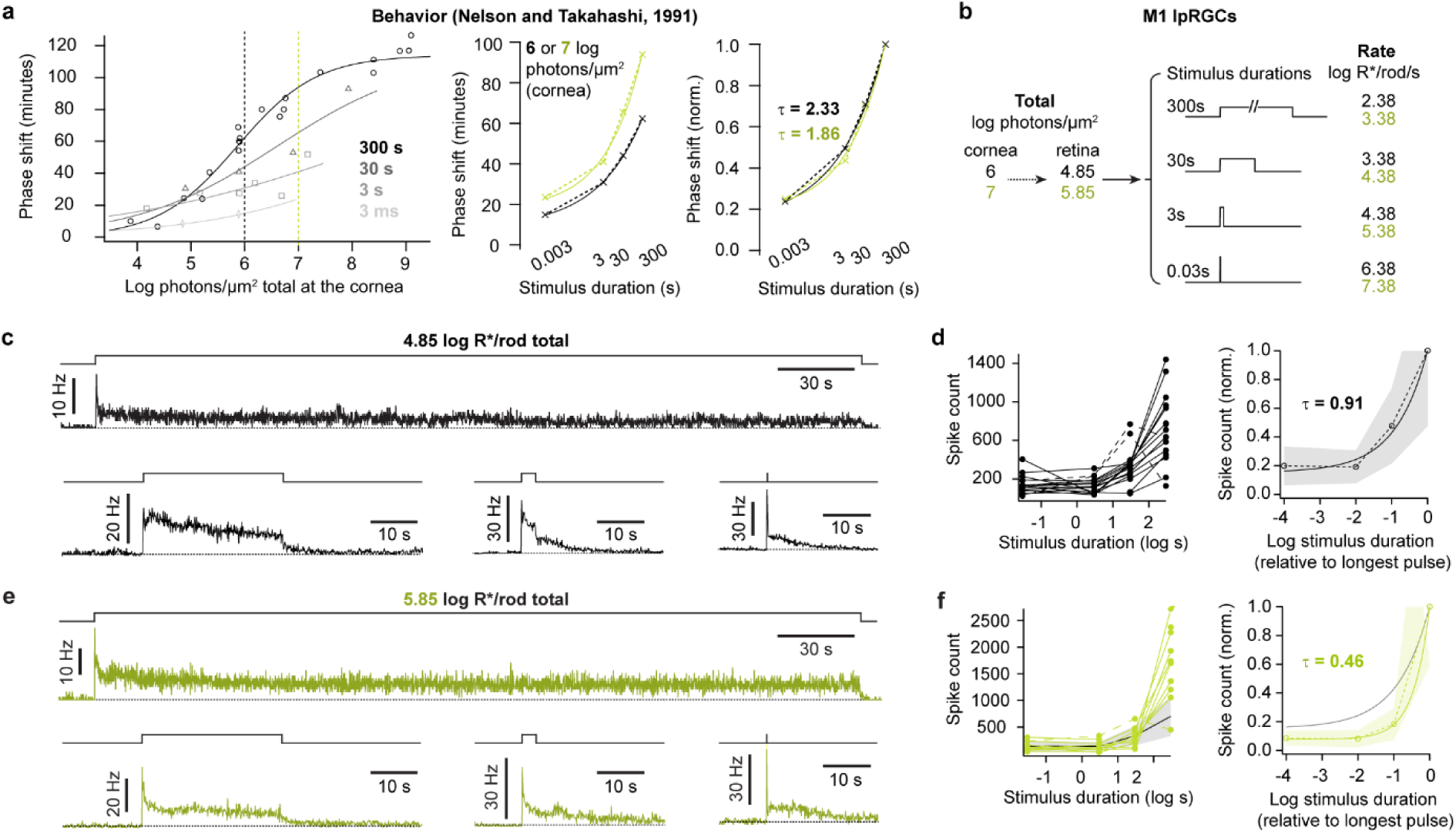
Precise and extended temporal integration by M1 ipRGCs. **a.** *Left,* Behavioral phase shifts defined by Nelson and Takahashi^5^ to various photon counts (per unit area) delivered over 300 s (circles), 30 s (triangles), 3 s (squares), or 3 ms (diamonds). Fits are modified Naka-Rushton functions^5^. *Center,* Duration-response (D-R) relations for phase shifting at two photon counts at the cornea: 6 (black) and 7 (green) log photons/μm^2^ (**Methods**). Each D-R relation is fit with a single exponential. *Right,* Normalized D-R relations with the time constants of their single exponential fits. **b.** The stimulus suite, modeled on those of **a**, used to examine the precision of temporal integration by M1s. Two photon counts are tested, which yield 4.85 (black) and 5.85 (green) log R*/rod at the retina (broadband light). These photon counts correspond to 6 and 7 log photons/μm^2^ at the cornea (**Methods**). **c.** Average spike firing rates of M1s in response to 4.85 log R*/rod delivered over 300 s (17 cells), 30 s (15), 3 s (15), and 30 ms (16). Stimulus monitor at top. Loose-patch recording, 35 °C, no synaptic antagonists, 0.1-s bins. **d.** *Left,* Duration-firing (D-F) relations for all M1s in **c**. Most relations were fit by a single exponential (τ = 0.86 ± 0.27 log units, 11 of 17 cells). Dashed traces indicate unusual, non-monotonic D-F relations. *Right,* D-F relations were normalized to maximum and averaged (solid line; shading is ±1 SD). The fit is a single exponential with τ = 0.91. **e.** As in **c**, for a different set of M1s (10 cells) given 5.85 log R*/rod over different durations. **f.** As in **d** but for the 5.85 log R*/rod stimulus suite. The single exponential fit to the average has τ = 0.46. For comparison, the average D-F (left) and single exponential fit to the average (right) are shown for the 4.85 log R*/rod stimulus suite.

We began with a stimulus producing 4.85 log R*/rod at the retina, which is near an intensity used behaviorally (6 log photons/µm^2^ at the cornea; **Fig. 3a-b**). Delivering this stimulus over different durations produced distinct spike patterns in M1s (**Fig. 3c**, **Extended Data Fig. 6**). Surprisingly, the total spike count had a shallow dependence on duration for most M1s. Their individual duration-firing (D-F) relations were well described by single exponentials with a mean time constant (τ) of 0.86 ± 0.27 log s, similar to the population τ of 0.91 log s (11 cells; **Fig. 3d**). In other words, across the 10^4^-fold span of tested durations, the spike count varied by only ∼5-fold. Variation was steepest from 30 to 300 s but was only ∼2.1-fold in this 10-fold range for the population. Thus, M1s exhibit high temporal integration. They count photons with precision over minutes.

Next, we considered a photon count producing 5.85 log R*/rod (**Fig. 3e**). These more intense stimuli evoked higher spike counts (**Fig. 3f**), indicating intensity encoding by M1s. The D-F relation was steeper at this higher intensity (**Fig. 3f**; population τ = 0.46 log s). Nevertheless, across the 10^4^-fold span of durations, the population spike count varied by only ∼12-fold; in the steep region from 30 to 300 s, variation was ∼5.5-fold.

Nelson and Takahashi also observed precise temporal integration of stimuli that differed in their temporal structure; for example, a given photon count received steadily or in trains of discrete pulses^6^. We compared M1 spiking for constant and pulsed stimuli that produced 4.85 or 5.85 log R*/rod in 300 s, a duration that is especially effective at driving behavioral phase shifts^5,6^. We gave pulses at 0.1 Hz and, in a subset, 1 Hz (**Fig. 4**). At each photon count, despite varying temporal dynamics, most cells showed little difference in total spike count among steady and pulsed stimuli (**Fig. 4b-c**). This was the case whether synaptic drive appeared low (a cell that failed to follow 1-Hz pulses) or high (cells showing large, fast transients at pulse onsets), indicating that diverse synaptic and intrinsic balances support precise temporal integration. The higher photon count also produced a higher population spike count across both steady and 0.1-Hz pulsed stimuli, again consistent with M1s encoding the overall light intensity.

**Fig. 4.**
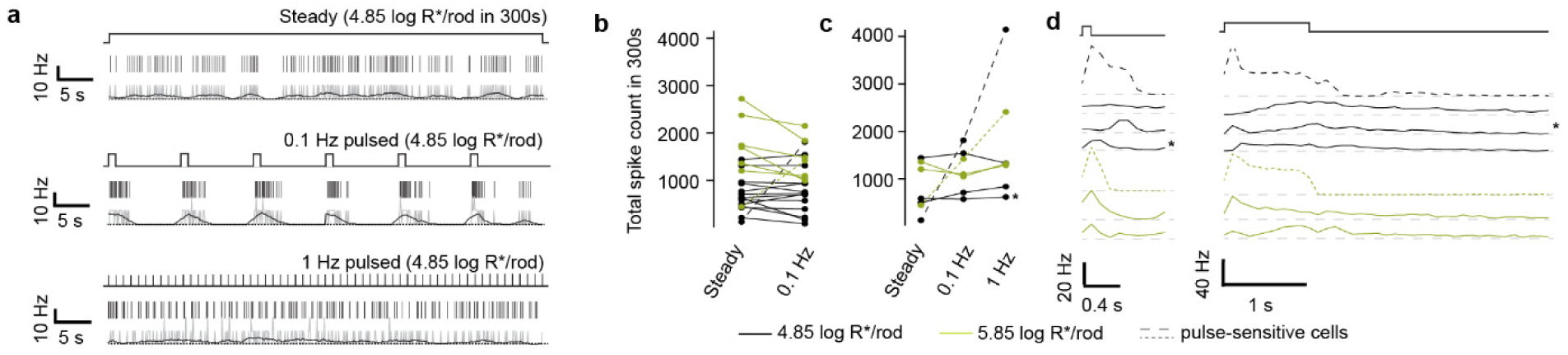
Precise integration of pulses by M1 ipRGCs. **a.** Spike firing of an example M1 during stimuli delivering 4.85 log R*/rod in 300 s. *Top,* 300-s steady step (2.38 log R*/rod/s). *Center,* 0.1-Hz pulses (1-s duration, 3.38 log R*/rod/s each). *Bottom,* 1-Hz pulses (0.1-s width, 3.38 log R*/rod/s each). Response excerpts are taken ∼90-150 s after stimulus onset, and each shows the stimulus (top), spike rasters (center), and spike rate histograms (0.1-s bins in gray and 0.5-s moving average in black). Loose-patch recording, 35 °C, no synaptic antagonists. **b.** Total M1 spike counts for the 300-s uniform step and 0.1-Hz pulsed stimulus, for 4.85 (black, 14 cells) or 5.85 (green, 7 cells) log R*/rod. Spike counts differed between photon counts (0.1 Hz, 4.85 vs. 5.85, p = 0.006; two-tailed Mann-Whitney U test). They did not differ detectably between uniform and pulsed presentations of those counts (4.85, uniform vs. 0.1 Hz, p = 0.90, and 5.85, uniform vs. 0.1 Hz, p = 0.30; paired Wilcoxon signed-rank test). **c.** A subset of 7 M1s received all three stimuli depicted in **a**, with total R* counts of 4.85 log R*/rod (black, 4 cells) or 5.85 log R*/rod (green, 3 cells). Two cells (dashed lines) preferred higher temporal frequencies, while most (solid lines) were insensitive to temporal structure. The example cell in **a** is marked with an asterisk. **d.** Pulse-triggered average spike rates of the 7 cells in **c**, given the 1- and 0.1-Hz stimuli (left and right, respectively; 0.1-s bins). For the 1-Hz stimulus, only the first 60 pulses are averaged to minimize any effect from adaptation. Shaded areas show the pulse timing. The example cell in **a** is marked with an asterisk.

### Intrinsic and synaptic origins of temporal integration by M1 ipRGCs

We recorded from M1s in synaptic antagonists and gave light in a continuous pulse producing 4.85 log R*/rod over different durations (**Fig. 5**). Transient spiking at light onset was smaller and slower than in physiological conditions. Spike counts were also lower across all stimulus durations. Despite these effects of synaptic antagonism, the shape of the population D-F curve was essentially unchanged (**Fig. 5b**). The population τ was 0.93 with antagonists and 0.91 without. In both conditions, across the 10^4^-fold range of durations, the population spike count varied by ∼5-fold; between 30 and 300 s, it varied by ∼2-fold. Hence, synaptic inputs largely result in a scaling of the population spike count set by intrinsic activity—despite the different sensitivities and kinetics of these two drives—supporting precise temporal integration. One exception to this scaling is that, at the end of the 300-s step, the population spike rates without and with synaptic antagonists became largely indistinguishable (**Fig. 5a**). Over long time scales, temporal integration relies largely on melanopsin phototransduction.

**Fig. 5.**
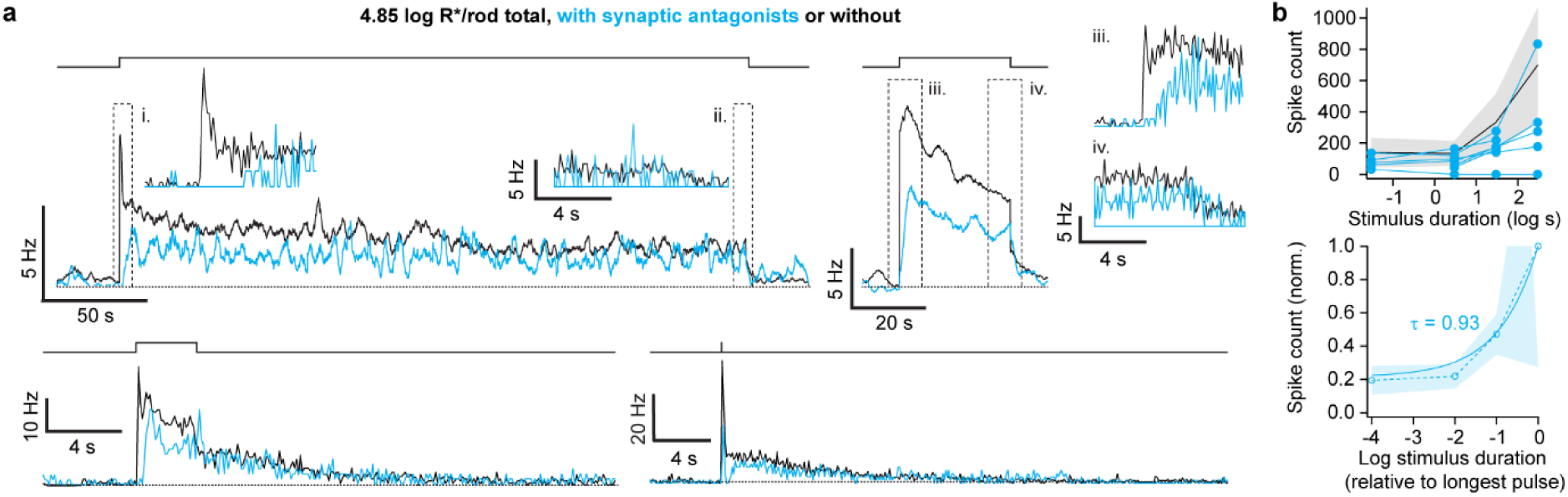
Synaptic input supports the temporal integration of M1 ipRGCs. **a.** Average spike firing rates of M1s given 4.85 log R*/rod over 300, 30, 3, and 0.03 s. Cyan: Synaptic antagonists present (5-8 cells per stimulus duration). Black: Same data as in Fig. 3c. Firing rates are shown as 0.5-s moving averages for clarity, and insets are binned at 0.1 s to resolve transients. Loose-patch recording, 35 °C. **b.** *Top,* Individual D-F relations of the M1s in synaptic antagonists (cyan, 9 cells). The average D-F for M1s without antagonists is provided for comparison (mean and SD in black and gray, respectively, same data as in Fig. 3d). *Bottom,* Population average of D-F relations of M1s in synaptic antagonists. Single exponential fit, τ = 0.93 (solid line). Shading is ±1 SD.

### Pupil constriction raises the temporal integration of M1 ipRGCs to behavioral precision

We hypothesized that temporal integration is more precise *in vivo* because longer stimuli are attenuated by pupillary constriction, flattening the D-F relation. To test this idea, we measured the pupil of behaving mice while giving light producing 4.85 log R*/rod over different durations (**Fig. 6a**). Pupillary constriction was undetectable during the briefest stimulus. For the other three durations, we measured the time courses of constriction and designed stimuli that mimic their attenuations of light intensity (**Fig. 6b**). We delivered these pupil-corrected stimuli to M1s in the *ex vivo* retina under physiological conditions. Temporal integration was more effective for these stimuli (population D-F relation τ of 2.37 log s, 7 cells; **Fig. 6c-d**). Across the 10^4^-fold difference in stimulus durations, the population spike count varied by ∼3.8-fold; from 30 to 300 s, it varied by 1.4-fold. The D-F relation closely matched that of behavioral phase shifting measured by Nelson and Takahashi (**Fig. 6d**). We conclude that pupil constriction increases the precision of circadian photoregulation by tuning the temporal integration of M1s.

**Fig. 6.**
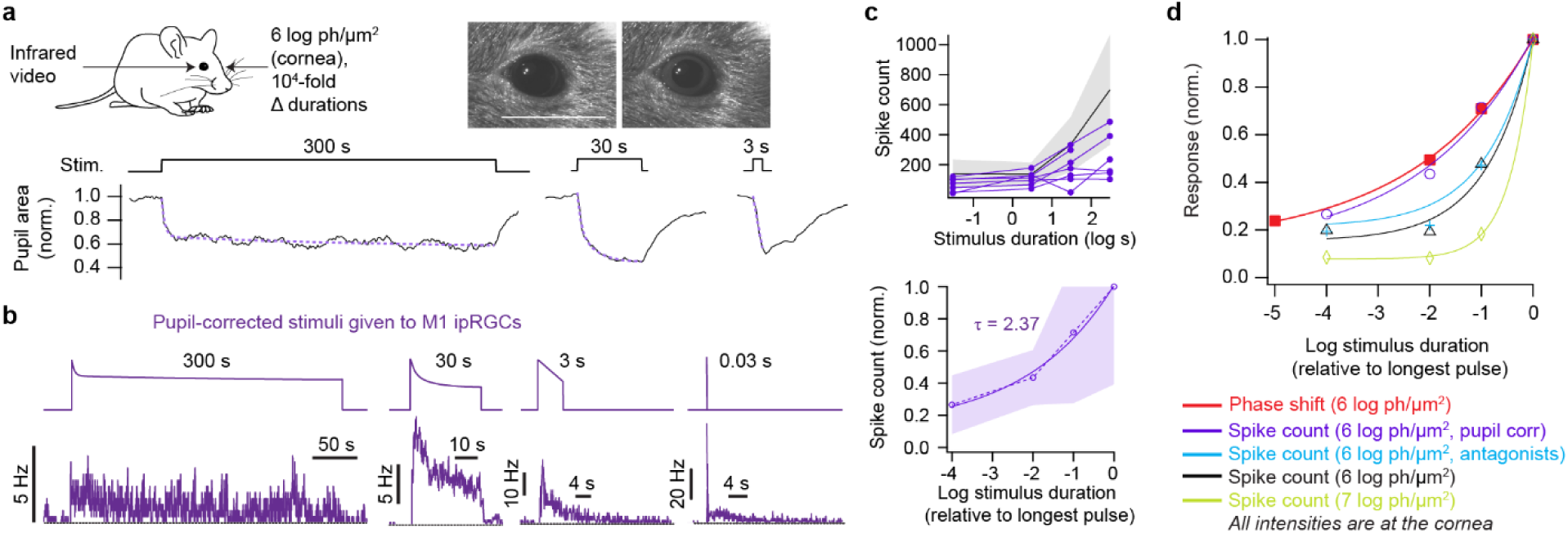
Pupillary constriction tunes the temporal integration of M1 ipRGCs. **a.** *Top,* Pupillometry during delivery of 4.85 log R*/rod stimuli (**Methods**). Individual video frames show the pupil of an example mouse in darkness (left) and at the peak of its response to a 300-s light pulse (right). Scale bar: 5 mm. *Bottom,* Pupil diameter, normalized to its dark-adapted level, for three stimulus durations (black; average of 8 trials for 4 mice). Exponential fits (double: 300 s, single: 30 s/3 s) are overlaid in violet. **b.** Light pulses (top) were attenuated according to pupil constriction to make pupil-corrected stimuli. Average spike rates are given (bottom; 6-7 cells per stimulus condition) for each pupil-corrected stimulus (300, 30, and 3 s) and an uncorrected 30-ms stimulus (**Methods**). **c.** *Top,* D-F relations for single M1s in **b** (violet, 7 cells). The average D-F from the pupil-uncorrected, 4.85 log R*/rod group is shown for comparison (black and gray; same data as in Fig. 3d). *Bottom,* D-F relations of the pupil-corrected group were normalized to their maxima and averaged (dashed line). Single exponential fit (solid line), τ = 2.37. Shading is ±1 SD. **d.** Population D-F relations of all four experimental M1 groups. A behavioral phase-shifting D-R from Nelson and Takahashi is plotted for comparison (6 log photons/μm^2^, equivalent to the 4.85 log R*/rod given to M1s; see Fig. 3b). With pupil correction, temporal integration by M1s resembles that of animal behavior.

## DISCUSSION

M1 ipRGCs are essential for circadian regulation. They provide most retinal innervation of the brain’s principal clock, are obligate relays of rod and cone signals to this structure, and set the clock even with only their intrinsic photosensitivity^22,23^. We have investigated how the functional properties of M1s support this role.

We developed a bioluminescence method to identify these rare neurons without desensitizing the retina. This method facilitates investigation of cellular and circuit mechanisms in a behaviorally relevant context. We introduce a mouse line and viral vectors for reporter expression; the substrate is readily available and titratable; and suitable cameras are more economical than multiphoton fluorescence systems. Incorporating reporters of different spectra and higher emission could increase the accessibility and flexibility of the approach^24,25^.

M1s are understood to encode light intensity^9^. The apparent simplicity of the task contrasts with the complexity of M1 light responses at the level of single cells (e.g., adaptation, depolarization block, and heterogeneity) and of their microcircuitry^11,13,14,17,26–29^. We find that these responses account for two hallmarks of clock photoregulation observed in animal behavior^5,6^. The first is its dynamic range, with an elevated threshold permitting vision in dim light without circadian resetting. The most sensitive M1s account for this threshold; as light intensifies and they collectively saturate, other M1s are recruited. The second hallmark is the clock’s temporal integration, which counts photons with precision over minutes to promote the encoding of overall illumination, a proxy for solar elevation. Response complexity supports this process across orders-of-magnitude changes in environmental intensity. For example, higher light intensities produce more adaptation and depolarization block to trim spike outputs, and different intensities engage these mechanisms at distinct levels in varied subsets of M1s (**Extended Data Fig. 7**).

We find that pupil constriction tunes the M1 population such that its precision of temporal integration matches that observed behaviorally (**Fig. 6d**). Pupil constriction is stronger even than synaptic input in shaping the temporal integration of M1 outputs (**Fig. 6d**). The level of temporal integration in behavioral phase shifting is constant across a broad range of intensities^5^ (**Fig. 3a**). This suggests that pupillary constriction operates accurately over this range to support circadian photoregulation, as it does for visual perception^30^. Pupil constriction, like circadian photoregulation, is largely driven by M1s^9^. Hence, distinct processes influenced by these cells appear to be coordinated.

M1s are classified as a single cell type by physiological, morphological, and molecular parameters. Variations in these parameters support subdivisions of M1s, though these groups do not cohere across parameters^8,11,13,14,23,29,31,32^. In our sample, some act more as photoreceptors and others as relays for rod/cone signals. For those with identified locations, the former were enriched in the ventral retina (10/10 identified ventral cells were high-intrinsic, versus 3/5 in the dorsal retina) and the latter in the dorsal retina (2/5 identified dorsal cells were high-synaptic, versus 0/10 in the ventral retina), consistent with dorsal M1s showing lower melanopsin immunoreactivity and higher dendritic complexity^14,29,31^. Future studies may provide a logic to these variations. They may also determine if our high-synaptic and high-intrinsic grouping is more than an operational definition, which we made to facilitate investigation of how cellular heterogeneity supports behavior.

A caveat of our study is that temporal integration of circadian phase shifting was defined using hamsters^5,6^, while our experiments required expanding upon tools established over decades for mice. While species comparisons require care, we found exquisite agreement between the two data sets. This concern is also mitigated by the conservation of M1 signaling mechanisms between mice and an even more distant species, the macaque monkey^12^.

Neurons encode salient aspects of the environment. We find that the first steps in photoreception select for indicators of Earth’s rotational phase. That is, M1 neurons of the retina produce highly customized information for clock regulation. Performing these computations at this early stage makes them available to the dozens of brain regions that M1s innervate^33^. Thus, diverse aspects of the nervous system may receive cues of the day-night cycle.

## ACKNOWLEDGEMENTS

The authors thank Todd Anthony for generating nano-lantern expression constructs; Dwight Nelson and Joseph Takahashi for sharing raw data; Alan Emanuel, Alex Chen, and Wendy Liu for preliminary experiments; Mantu Bhaumik for mouse genetic engineering; Siva Nagappan and Hisashi Umemori for guidance in molecular genetics; Marie Burns for permission to use *Gnat2^−/−^* mice; King-Wai Yau for *Gnat2^−/−^* and *Opn4:Cre* BAC transgenic mice; Vladimir Kefalov for *Gnat1^−/−^* mice; Samer Hattar for *Opn4^Cre^* knock-in mice; Chen Wang, Yiming Zhou, and Zhigang He for virus production; and Tiago Branco for support. Funding was provided by National Institutes of Health grants EY025466 to ESM; EY033639 and EY037374 to FSC-H; 1U54HD090255 to Boston Children’s Hospital IDDRC; P30 EY012196 to Harvard Medical School; and EY023648, EY034089, EY025555, EY032731, and EY036071 to MTHD. Funding was also provided by grants from the Alcon Research Institute, Karl Kirchgessner Foundation, Knights Templar Eye Foundation, March of Dimes, and Whitehall Foundation to MTHD.

## AUTHOR CONTRIBUTIONS

ESM: Conception, experiment design, execution, analysis, interpretation, and writing. HAB: Electrophysiology of α-RGCs and pupillometry. NM: Genetic characterization of the NL mouse line. MCB: Analysis. FC-H: Retrograde labeling and histology. N-MV: Retrograde labeling. TN: Provision of NL constructs. MTHD: Conception, experiment design, interpretation, and writing.

## DATA AND CODE AVAILABILITY

Data and code are available upon request.

---

**Extended Data Table 1.** Minimal desensitization with bioluminescence imaging. Mean ± 1 SD reported for each condition. Statistically significant comparisons (following Bonferroni correction) are italicized.

|  | **Baseline Firing**<br>**(Hz)** | **p** | **Flash Threshold**<br>**(log<sub>10</sub> R\*/rod)** | **p** |
| --- | --- | --- | --- | --- |
| **NL<sup>+</sup> M4**<br>**ipRGC** | 9.2 ± 10.9 Hz (4<br>cells) |  | -1.70 ± 0.48 (4<br>cells) |  |
| **CH<sup>+</sup> M4**<br>**ipRGC** | 5.4 ± 8.6 Hz (4<br>cells) | **vs. NL<sup>+</sup> M4:**<br>0.99 | -0.88 ± 0.69 (4<br>cells) | **vs. NL<sup>+</sup> M4:**<br>0.086 |
| **NL<sup>+</sup>CH<sup>+</sup> M4**<br>**ipRGC** | **All time points:**<br>25.1 ± 26.3 Hz<br>(12 cells) | **>150 min vs.**<br>**CH<sup>+</sup> M4:** 0.89 | -1.04 ± 0.31 (12<br>cells) | **vs. CH<sup>+</sup> M4:**<br>0.933 |
|  | **>150 min:** 4.5 ±<br>6.5 Hz (6 cells) | **>150 min vs.**<br>**NL<sup>+</sup> M4:** 0.91 |  | **vs. NL<sup>+</sup> M4:**<br>*0.013* |
| **OFF**<br>**sustained α-**<br>**RGC (in NL<sup>+</sup>**<br>**mice)** | 15.5 ± 6.8 Hz (6<br>cells) | **vs. NL<sup>+</sup> M4:**<br>0.35 | -1.98 ± 0.69 (6<br>cells) | **vs. NL<sup>+</sup> M4:**<br>0.686 |
|  |  | **vs. CH<sup>+</sup> M4:**<br>0.11 |  | **vs. CH<sup>+</sup> M4:**<br>0.028 |
|  |  | **vs. NL<sup>+</sup> CH<sup>+</sup>**<br>**M4:** 0.82 |  | **vs. NL<sup>+</sup> CH<sup>+</sup>**<br>**M4:** *0.008* |
| **NL<sup>+</sup>CH<sup>+</sup> M1**<br>**ipRGC** | 2.8 ± 3.3 Hz (32<br>cells) | **vs. NL<sup>+</sup> CH<sup>+</sup> M4**<br>**(>150 min):**<br>0.043 | 0.50 ± 0.73 (32<br>cells) | **vs. NL<sup>+</sup> CH<sup>+</sup>**<br>**M4:** *0.001* |

**Extended Data Table 2.**
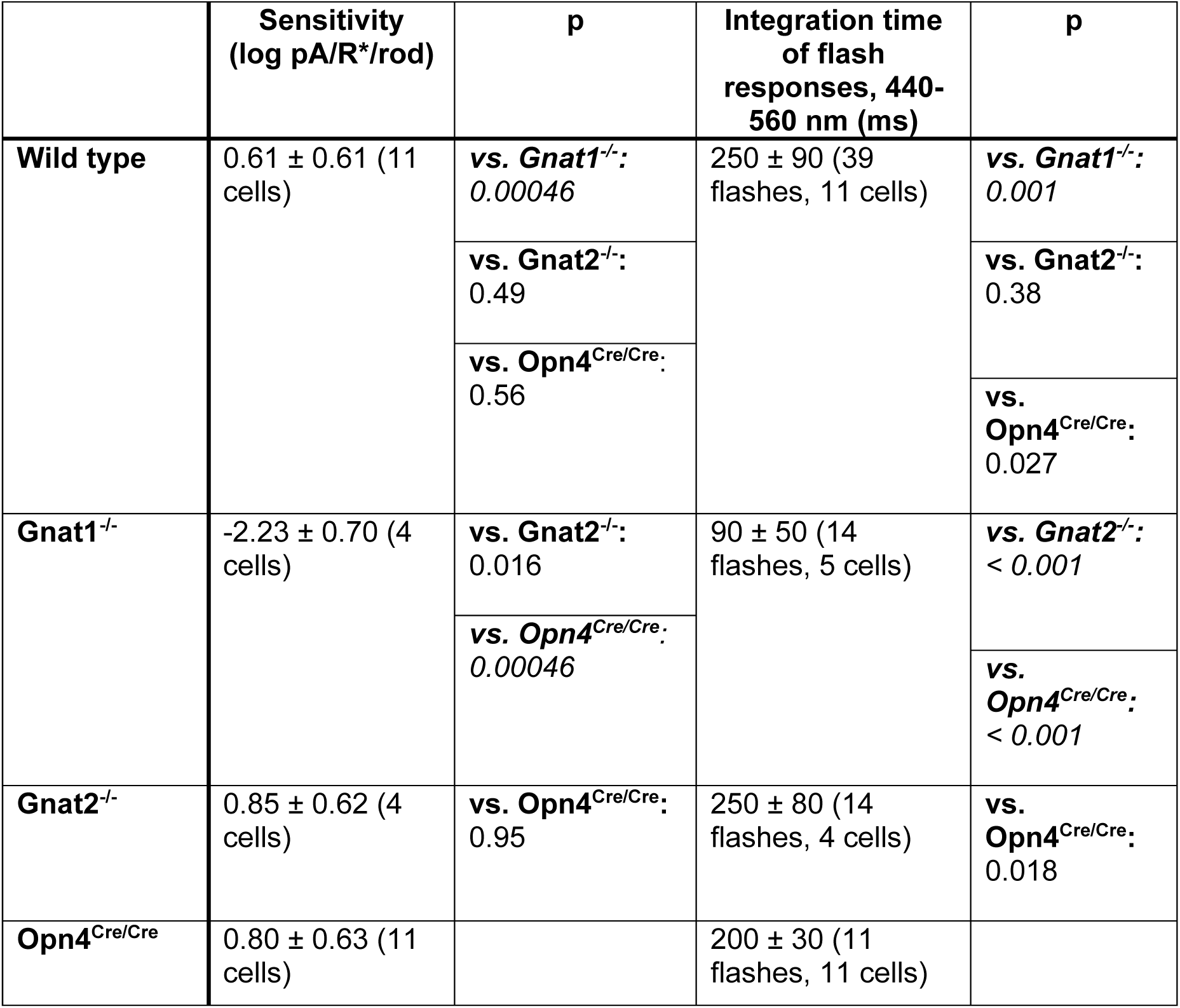
Sensitivity and kinetics indicate rod origins of threshold responses in M1 ipRGCs. Statistically significant comparisons (following Bonferroni correction) are italicized.

|  | Sensitivity<br>(log pA/R*/rod) | p | Integration time<br>of flash<br>responses, 440-560 nm (ms) | p |
| --- | --- | --- | --- | --- |
| **Wild type** | 0.61 ± 0.61 (11<br>cells) | ***vs. Gnat1<sup>-/-</sup>:***<br>*0.00046* | 250 ± 90 (39<br>flashes, 11 cells) | ***vs. Gnat1<sup>-/-</sup>:***<br>*0.001* |
|  |  | ***vs. Gnat2<sup>-/-</sup>:***<br>0.49 |  | ***vs. Gnat2<sup>-/-</sup>:***<br>0.38 |
|  |  | ***vs. Opn4<sup>Cre/Cre</sup>:***<br>0.56 |  | ***vs.***<br>***Opn4<sup>Cre/Cre</sup>:***<br>0.027 |
| **Gnat1<sup>-/-</sup>** | -2.23 ± 0.70 (4<br>cells) | ***vs. Gnat2<sup>-/-</sup>:***<br>0.016 | 90 ± 50 (14<br>flashes, 5 cells) | ***vs. Gnat2<sup>-/-</sup>:***<br>*< 0.001* |
|  |  | ***vs. Opn4<sup>Cre/Cre</sup>:***<br>*0.00046* |  | ***vs.***<br>***Opn4<sup>Cre/Cre</sup>:***<br>*< 0.001* |
| **Gnat2<sup>-/-</sup>** | 0.85 ± 0.62 (4<br>cells) | ***vs. Opn4<sup>Cre/Cre</sup>:***<br>0.95 | 250 ± 80 (14<br>flashes, 4 cells) | ***vs.***<br>***Opn4<sup>Cre/Cre</sup>:***<br>0.018 |
| **Opn4<sup>Cre/Cre</sup>** | 0.80 ± 0.63 (11<br>cells) |  | 200 ± 30 (11<br>flashes, 11 cells) |  |

**Extended Data Table 3.**
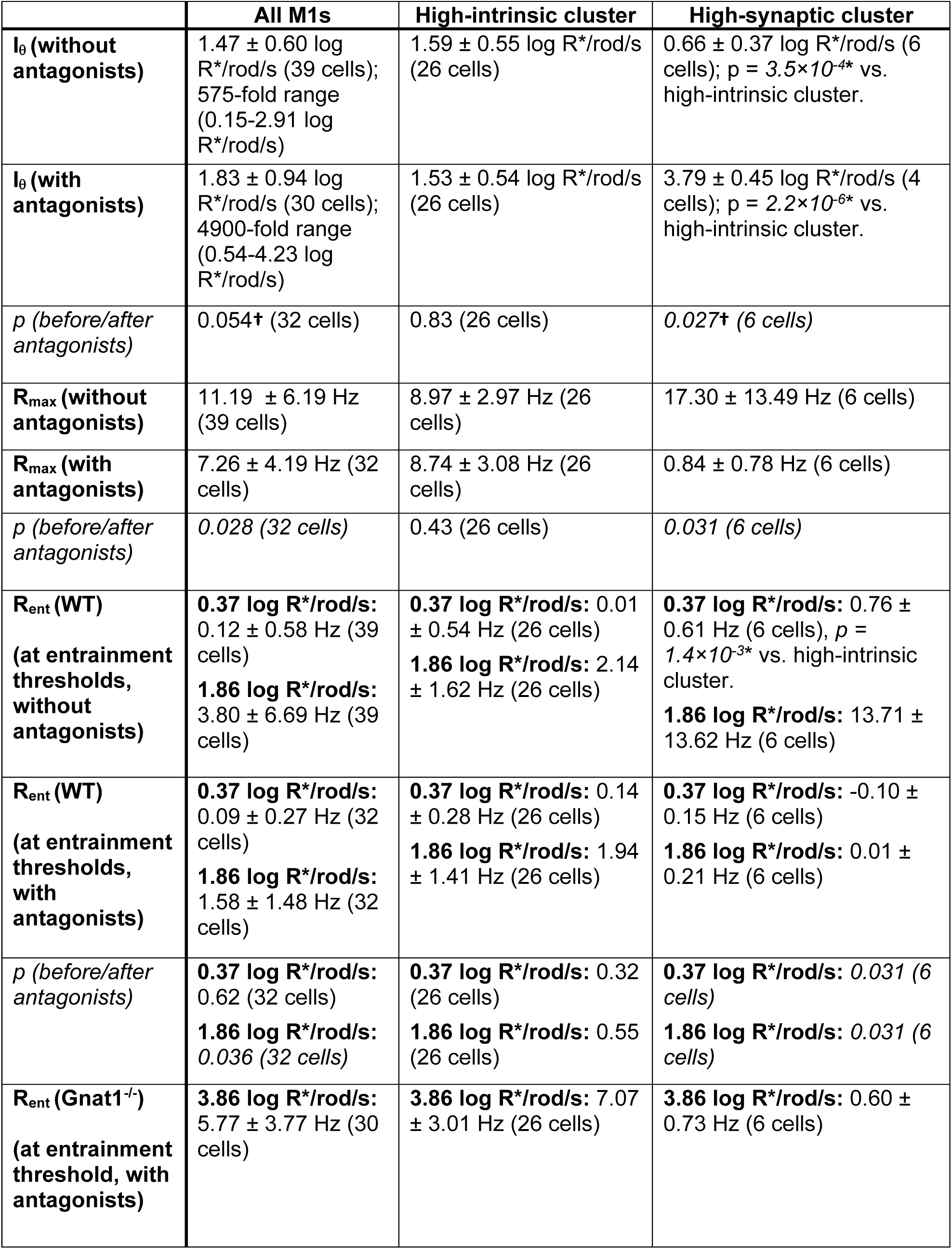
Varying balance of intrinsic and synaptic drives across M1s. Mean ± 1 SD reported for each condition. All firing rate values are corrected for baseline rates in darkness. I_θ_: threshold intensity at which steady firing rate increases by 1 Hz over dark value. R_max_: maximum steady firing rate across all staircase intensities. R_ent_: steady firing rate at published threshold intensities for circadian photoentrainment: 0.37 log R*/rod/s (50% of animals entrained) and 1.86 log R*/rod/s (80% of animals entrained) ^2,3^. Significance testing was performed using the paired Wilcoxon signed-rank test, two-tailed Mann-Whitney U test (*), or paired Prentice-Wilcoxon test (**†**) when appropriate (**Methods**).

**Extended Data Fig. 1.**
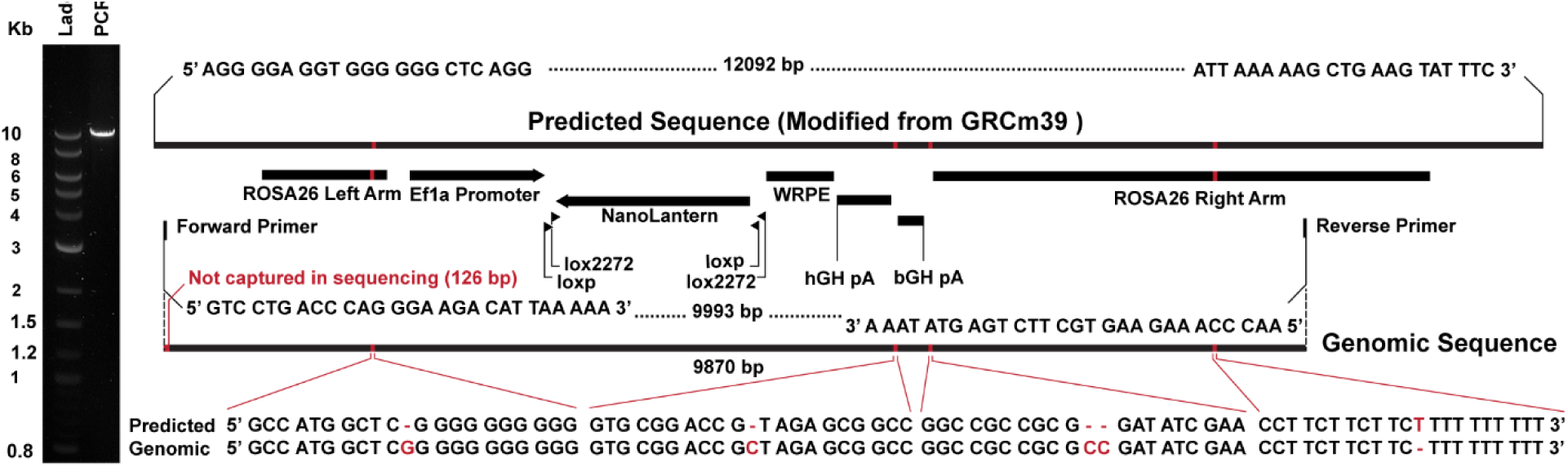
Validation of nano-lantern knock-in at the *ROSA26* locus, related to Fig. 1. *Left*, PCR product from the *ROSA26* locus of putative *ROSA26^nano-lantern/nano-lantern^* animals. *Right,* Alignment of the genomic sequence, derived from sequencing the purified PCR amplicon, against the mouse reference genome (GRCm39) modified to include the nano-lantern knock-in sequence. Solid horizontal lines indicate sequence alignments (black) and misalignments (red; not to scale). Sequence annotations, PCR primers and their annealing loci, as well as a 126-bp sequence not captured in sequencing are depicted between the predicted and genomic sequences.

**Extended Data Fig. 2.**
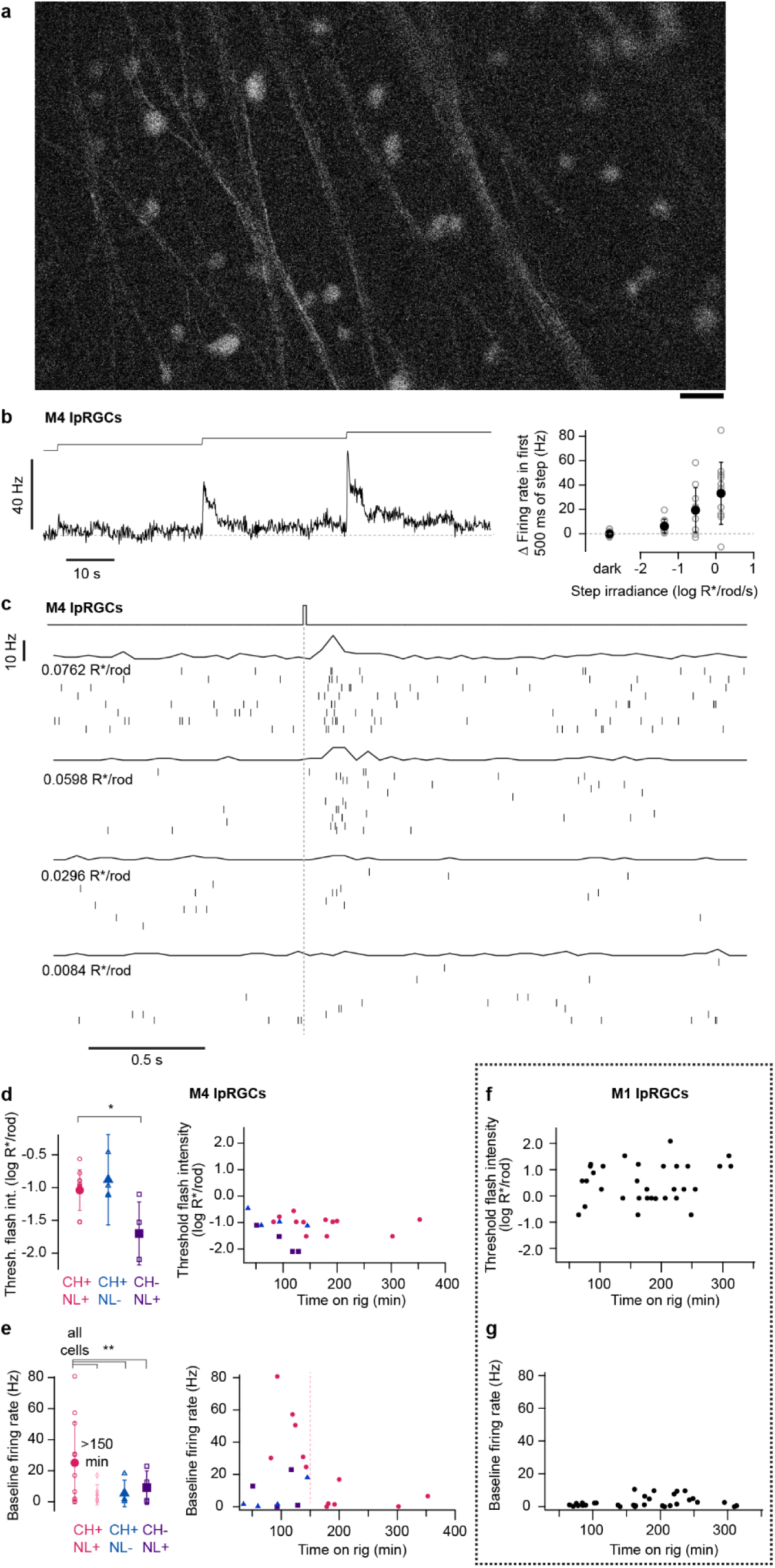
Bioluminescence imaging preserves the sensitivity of retinal ganglion cells, related to Fig. 1. **a.** Example of bioluminescence from an Opn4^Cre/+^;ROSA26^NL/+^ retina that was exposed to coelenterazine h (25 µM CH, 5 min) and then imaged (10-s exposure, CMOS camera). 20-µm scale bar. **b.** *Left,* M4 ipRGCs were identified by bioluminescence and their light responses (**Methods**). Spikes were recorded in loose-patch mode at 35 °C. A staircase was given from -1.4 to 0.1 log R*/rod/s. The average firing rate of 11 cells is shown (100-ms bins). Baseline firing in darkness is indicated (dashed line). The stimulus fully covered the dendritic arbor. *Right,* Transient increases in firing rate with increasing step intensity for the sample. Filled circles are the average ±1 SD. **c.** M4s were identified by bioluminescence and recorded as in **b**. Flashes of different intensities (15-30 ms, stimulus monitor at top, 410 nm) were given to estimate threshold. Spike firing rates (100-ms bins) are shown above sample rasters. The stimulus fully covered the dendritic arbor. **d.** *Left,* Summary statistics of flash threshold (the lowest flash intensity producing detectable spike modulation) of M4s (see **Extended Data Table 1** for statistics). Experimental cells were bioluminescent; they expressed nano-lantern (NL) and were exposed to CH (CH^+^ NL^+^, circles). Control cells were exposed to CH but did not express NL (CH^+^ NL^−^, triangles), or expressed NL but were not exposed to CH (CH^−^ NL^+^, squares). Test M4s had flash-driven thresholds that were comparable to M4s that lacked NL and CH, suggesting that these molecules do not greatly affect physiology. The difference between CH+ and CH-may be due to luminescence from CH oxidation^34^. *Right,* Flash thresholds plotted against time on the rig (which initiates CH washout and thus bioluminescence decay). M4 thresholds were stable over time. Spikes were recorded in loose-patch mode at 35 °C. **e.** As in **d**, but for baseline firing rate rather than threshold flash intensity. M4s in CH^+^ NL^+^ retinas showed elevated baseline firing initially (<150 min on the rig) but those recorded >150 min after CH washout (*right,* diamonds) had baseline firing rates that were indistinguishable from control M4s (see **Extended Data Table 1** for statistics). **f.** As in the right panel of **d**, but for NL-expressing M1 ipRGCs (transduced retrogradely from the hypothalamus). M1 thresholds show no dependence on time since washout; their thresholds are also higher than those of M4s (see **Extended Data Table 1** for statistics). **g.** As in **f**, but for baseline firing rather than threshold flash intensity. M1 firing rates in darkness showed no dependence on time since CH washout (see **Extended Data Table 1** for statistics).

**Extended Data Fig. 3.**
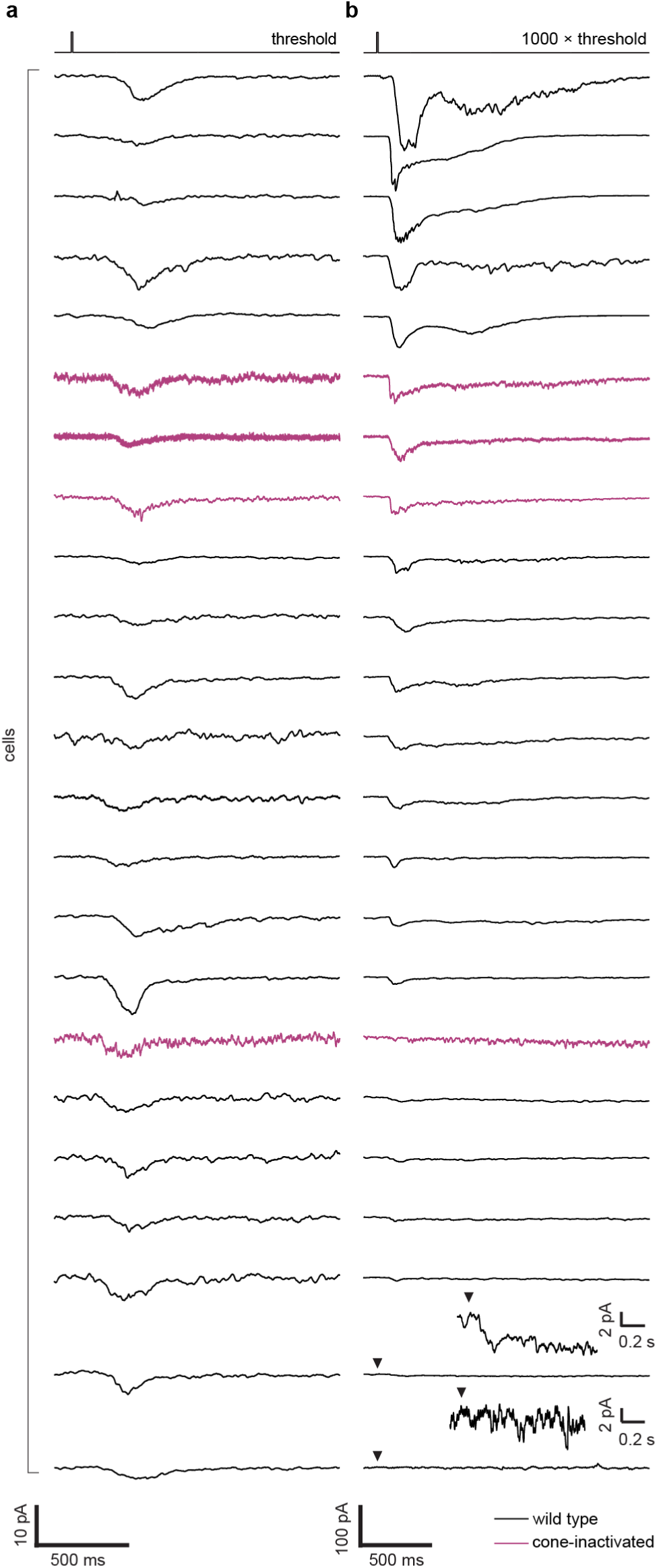
High flash intensities reveal the diversity of synaptic responses in M1 ipRGCs, related to Fig. 1. **a.** Near-threshold flash responses of 23 M1s, recorded in voltage clamp at -80 mV. Stimulus flashes (top trace showing square pulse) were 20-30 ms and 500 nm, and their intensity varied from 0.034-0.42 log R*/rod; response amplitudes varied from 2-10 pA. Cells are from wild-type (black) or *Gnat2^−/−^* (magenta) retinas. The latter have inactivated cones but normal rod and melanopsin responses. Traces are averages of 6-40 sweeps. Recordings were made at 23 °C for additional stability. **b.** Responses from these same 23 cells to flashes calibrated to 1000× each cell’s approximate threshold. Traces are averages of 1-5 sweeps. Cells in **a** and **b** are ordered by the strength of their response to this flash; response amplitudes varied from undetectable (bottom row) to ∼200 pA (top row). Insets show the two smallest responses (bottom two rows) on an expanded current base (arrowheads mark the onset of the flash).

**Extended Data Fig. 4.**
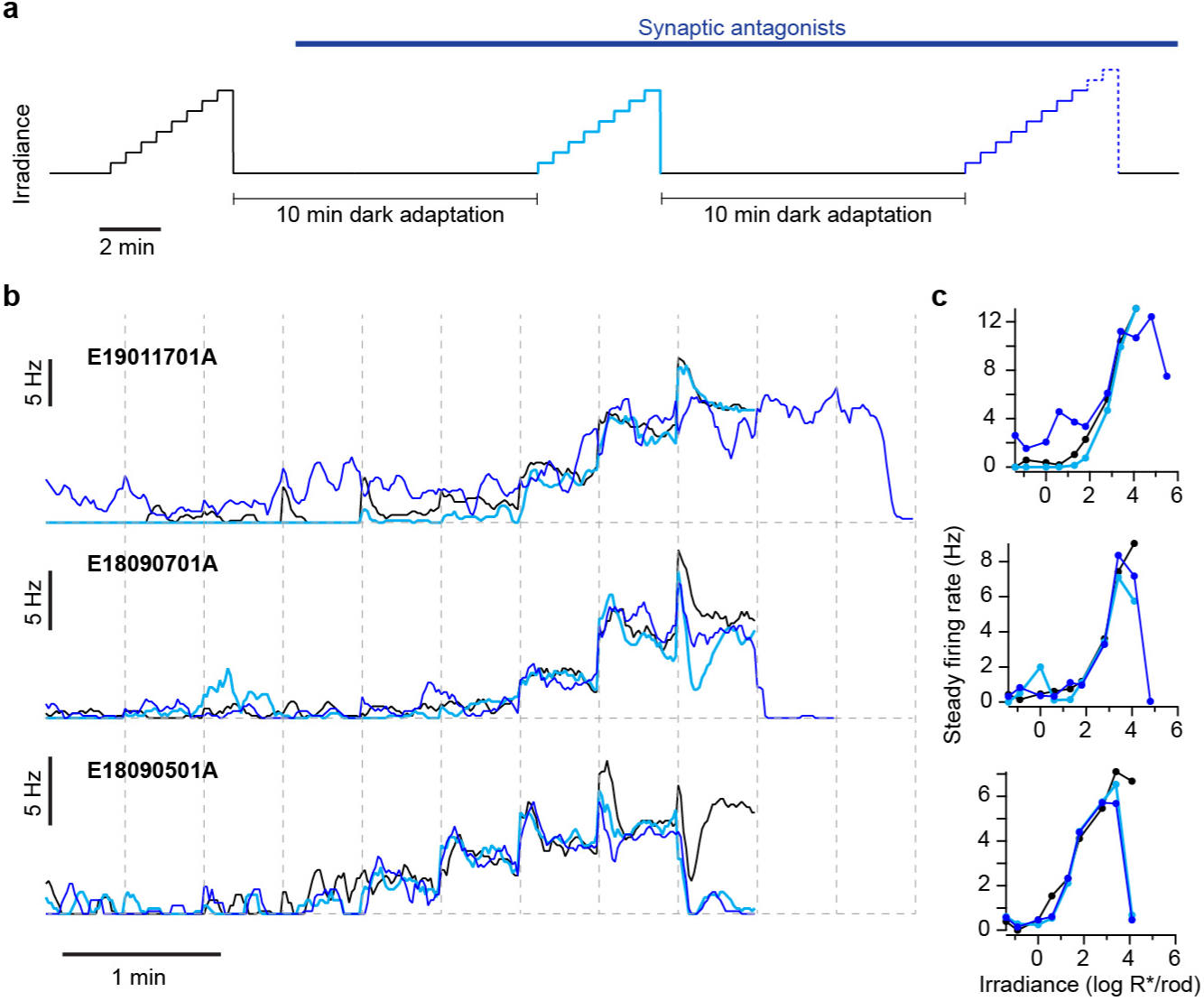
Stationarity of M1 ipRGC intrinsic photosensitivity during intensity staircases, related to Fig. 2. **a.** Cells were presented with an intensity staircase (black; 8 steps, 30-s each, -0.9 to 4.1 log R*/rod/s, 460-nm light). During the subsequent 10 minutes of dark adaptation, synaptic antagonists were added to isolate the intrinsic light responses (**Methods**). Antagonists remained throughout the rest of the recording. The same staircase was presented a second time (cyan) and, after another 10-min dark adaptation, a third time (blue). Occasionally, the third staircase included 1-2 additional intensity steps, which reached a maximum of 5.5 log R*/rod/s (dashed line). Recordings were performed in loose-patch mode at 35 °C. **b.** Overlaid firing rates from the three staircases (black, cyan, and blue traces) for three example cells (top, middle, and bottom rows; 5-s moving average). The last two epochs correspond to additional intensity steps (see **a**). **c.** Intensity-firing relations for the data in **b**.

**Extended Data Fig. 5.**
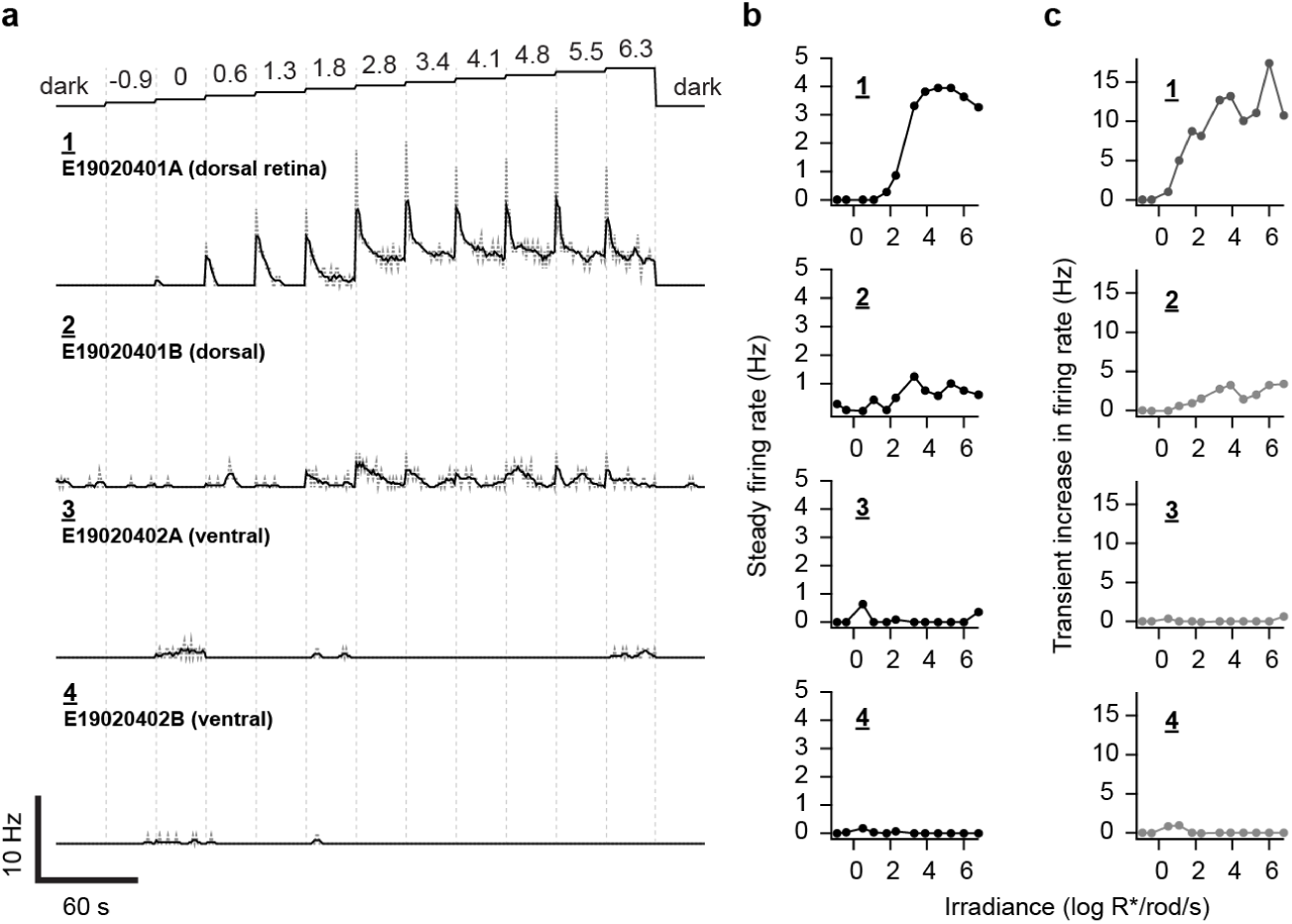
M1 ipRGC intensity-firing relations without melanopsin, related to Fig. 2. **a.** The extended staircase stimulus (see Fig. 2; all intensity values are in log R*/rod/s) was delivered to 4 Opn4^Cre/Cre^ M1s (rows 1-4). Cells 1 and 2 (**E19020401A/B**) were recorded simultaneously in the dorsal retina, and cells 3 and 4 (**E19020402A/B**) were recorded simultaneously in the ventral half of the same retina. Loose-patch recordings, no synaptic antagonists, 35 °C. **b-c.** Intensity-firing relations for the four cells in **a**, reflecting steady firing during the last 10 s of each step (**b**) or transient firing during the first 500 ms of each step (**c**). Cells 3 and 4 have virtually no light responses; cell 2 has a modest response with little intensity encoding, and cell 1 is able to encode intensity across several log units.

**Extended Data Fig. 6.**
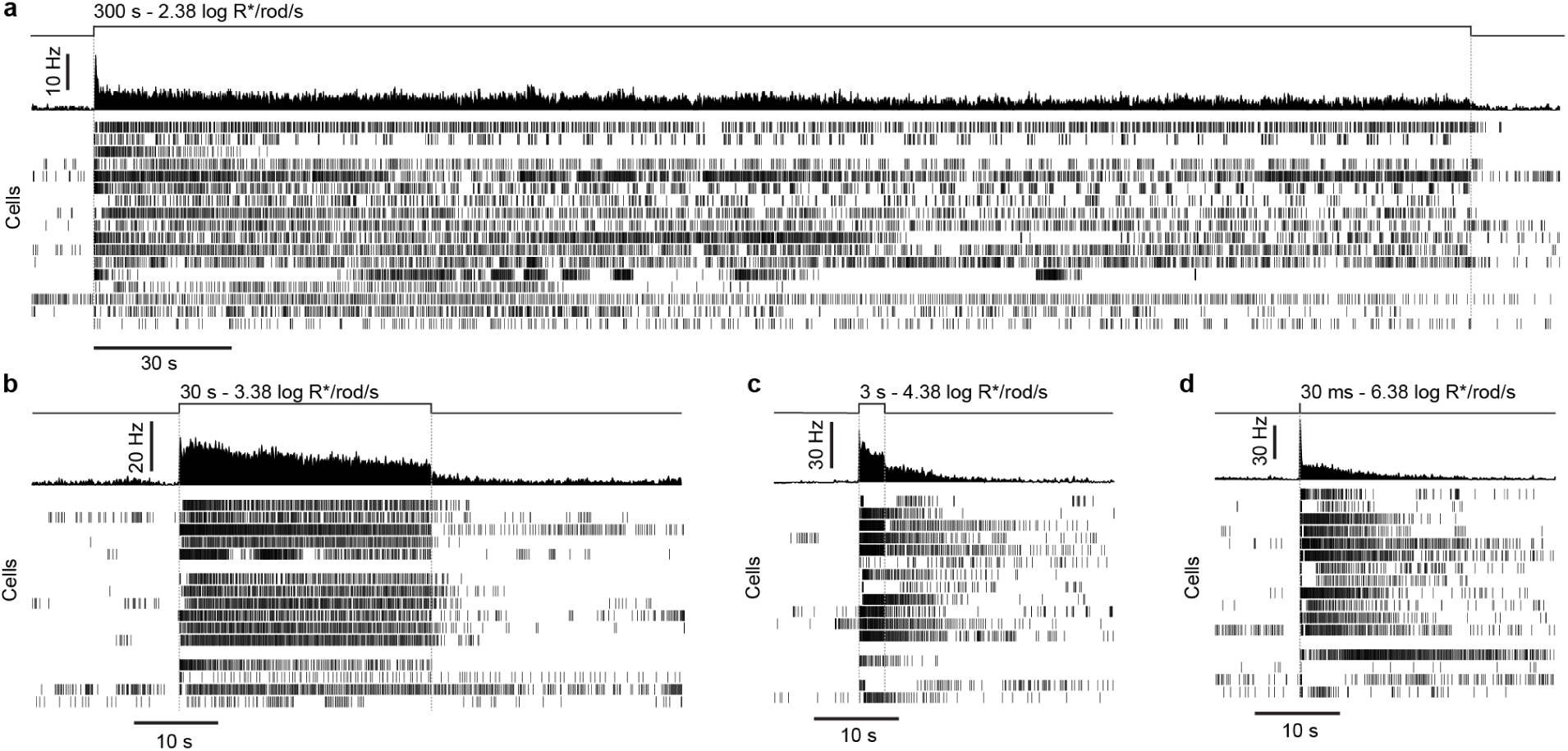
Individual M1 ipRGC responses to single-pulse temporal integration stimuli, related to Fig. 3. **a**-**d**. Average spike rates (top) and individual spike rasters for all M1 responses to the 300 s (**a**, 17 cells), 30 s (**b**, 15), 3 s (**c**, 15), and 30 ms (**d**, 16) stimuli. Each stimulus delivered a total of 4.85 log R*/rod. Stimulus monitors at top. Stimuli were delivered in random orders across cells, with >300 s of dark adaptation between presentations. Each row is one trial from one cell. Not all cells received all four stimulus durations; blank rows indicate these missing data. Loose-patch recording, 35 °C, no synaptic antagonists.

**Extended Data Fig. 7.**
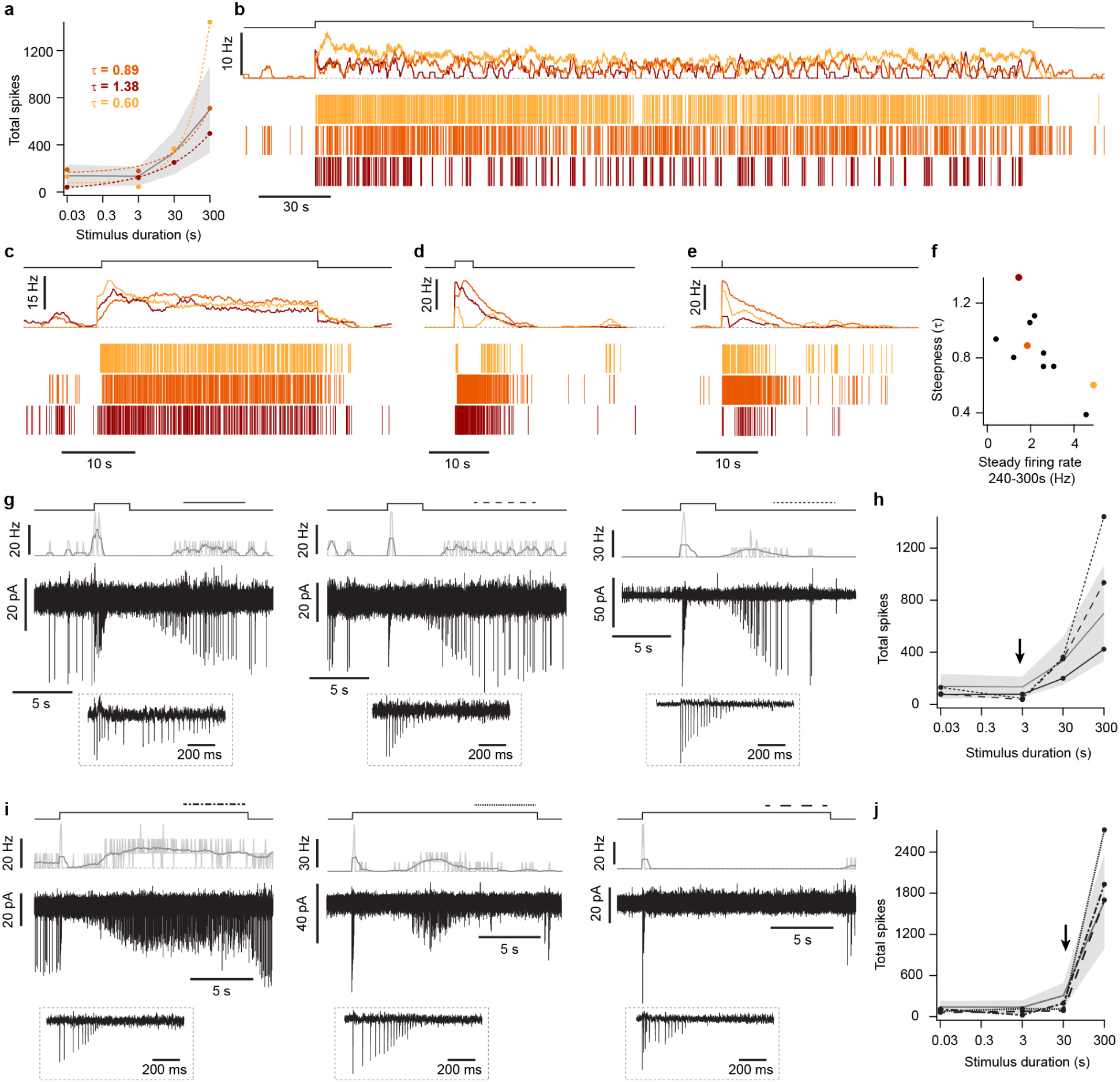
Sources of variability in temporal integration across M1 ipRGCs, related to Fig. 3. **a**. Duration-Firing (D-F) relations for three example M1s from the data set shown in Fig. 3d and **Extended Data Fig. 6**. Stimuli delivered a total of 4.85 log R*/rod over different durations, and produced varying spike numbers across the example cells. Individual data points (circles) are colored by cell identity and overlaid with single exponential fits (dashed lines, with τ reported for each). The black line and gray shading are mean ± SD. Loose-patch recording, 35 °C, no synaptic antagonists. **b-e**. Spike rates (above) and rasters (below) for the three example cells in **a** when given the 300 s (**b**), 30 s (**c**), 3 s (**d**), or 30 ms (**e**) stimulus. Histograms are shown as a 0.5-s moving average of 0.1-s bins. Stimulus monitors at top. **f.** The τ of all D-F relations in the data set of Fig. 3d (11 of 17 cells whose relations could be fit with a single exponential). Steeper D-F relations, which have less precise temporal integration, have lower τ values. τ was correlated with the steady firing rate (measured in the last 60 s of the 300-s stimulus; Spearman’s r = -0.68, p = 0.03). **g.** 3 example M1s for which the 3-s, single-pulse stimulus (4.85 log R*/rod) produces spike silencing. Shown are stimulus monitors (shaded), spike rates (light gray: 0.1-s bins, dark gray: 0.5-s moving average), and raw traces. Insets show the first 1 s after stimulus onset. Spike amplitude dwindling into noise, then recovering after stimulus cessation, is a signature of depolarization block^11^. Thus, depolarization block shapes temporal integration of M1s. **h.** D-F relations for the 3 cells in **g** (black lines with markers), with the population mean D-F relation for 4.85 log R*/rod (solid gray line, shading is ±1 SD) given for comparison. The arrow marks the stimulus that produced depolarization block in these cells. Note that all 3 have below-average responses to this stimulus. **i.** 3 example M1s given a 30-s stimulus delivering 5.85 log R*/rod (same photon flux density but 10-fold longer duration than for the cells in **g**; see Fig. 3e). Cells showed varying degrees of depolarization block: blocking transiently and resuming firing during the stimulus (*left*), transiently resuming firing mid-stimulus and blocking again (*center*), and staying in block throughout the stimulus and only resuming firing in darkness (*right*). **j.** D-F relations for the 3 cells in **i**, compared to the population mean D-F relation for 5.85 log R*/rod (solid green line, shading is ±1 SD). The arrow marks the stimulus that produced depolarization block in these cells. Note that all 3 have below-average responses to this stimulus.

## METHODS

### Animals

All animal procedures were approved by the Institutional Animal Care and Use Committee of Boston Children’s Hospital. Animals were maintained on a 12:12 light:dark cycle. Males and females were used. No overt variation with circadian time, sex, or age was observed.

### Generation of mice for Cre-dependent expression of nano-lantern from the ROSA26 locus (*ROSA26^FLEX-NL^*)

Mice were generated using a protocol for insertion of NL into the ROSA26 gene locus using CRISPR/Cas9^35^ (**Extended Data Fig. 1**). The guide RNA was 5’-ACTCCAGTCTTTCTAGAAGA-3’ (Millipore Sigma). To validate the line, a NL^+/+^ adult mouse was anesthetized and euthanized. ∼0.5 g of liver tissue was removed and digested overnight at 55 °C in 4 ml of 20 mM Tris (pH 8), 5 mM EDTA (pH 8), 400 mM NaCl, 1% SDS, and 400 µg/ml Proteinase K (Qiagen 19131). Digestion was quenched at 80 °C for 10 minutes. The sample was washed in buffer-saturated phenol (Thermo Fisher Scientific, 15513-39), phenol-chloroform-isoamyl alcohol (25:24:1, Thermo Fisher Scientific, 15593031), and twice in chloroform-isoamyl alcohol (24:1, Millipore Sigma, 25666; all volumes were 4 ml). For each wash, the sample was inverted (10×), incubated (10 min, ∼23 °C), and centrifuged (17,000 g, 1 min; room temperature, ∼23 °C); the aqueous phase was isolated for the next wash. DNA was precipitated by mixing the final aqueous phase with 4 ml of -20 °C ethanol and incubated for 40 min (-20 °C). DNA was transferred via a glass hook into 1.5 ml of 70% ethanol in double-distilled water (4 °C) and incubated on ice (15 min). The ethanol solution was decanted and the DNA was dried at 55 °C (∼20 minutes). It was then suspended in Buffer EB (10 mM Tris-Cl, pH 8.5, 500 µl; Qiagen 19086) via rocking overnight (4 °C). ∼23 µg of DNA was obtained (OD_260/280_ of 2.02 and OD_260/230_ of 2.16, measured using a NanoDrop Spectrophotometer; Thermo Fisher Scientific).

Primers flanking the *ROSA26* locus were designed using Primer Blast (NCBI) and manual optimization: Forward, 5’-GTCCTGACCCAGGGAAGACATTAAAAA-3’, and Reverse, 5’-AACCCAAAGAAGTGCTTCTGAGTATAAA-3’. Primers were acquired at 100 µM concentration in 10 mM Tris, pH 8.5 (Azenta). Four 50-µl PCR reactions, each including ∼200 ng of template DNA, were performed using PrimeSTAR GXL DNA polymerase (Takara, R050A), following manufacturer guidelines. PCR products were pooled, cleaned, and concentrated (Zymo, D4003) in 15 µl of Buffer EB. A ∼142 ng/µl product was obtained (OD_260/280_ of 1.88 and OD_260/230_ of 2.21). 1 µl of this product was diluted in 14 µl of nuclease-free water and mixed with 3 µl of 6X Gel Loading Dye (New England Biolabs, B7025S). The mixture was run on a 0.7% agarose gel with 1X SYBR Safe (Thermo Fisher Scientific, S33102) alongside a DNA ladder (New England Biolabs, N3200S), then imaged (Bio-Rad GelDoc XR). A single band was observed at ∼10 kb, matching the expected size of the PCR product (9,992 bp, **Extended Data Fig. 1**).

10 µl of the concentrated DNA product was submitted for long-read, end-to-end sequencing (Oxford Nanopore, Plasmidsaurus). Coverage was >100× per base. The vast majority of reads surpassed 9 kb in length. The consensus sequence matched that of the construct almost entirely. The only deviations within the knock-in sequence were outside of open reading frames and regulatory elements. The remaining misalignments are likely variations in the mouse genetic background (**Extended Data Fig. 1**).

### Mouse lines

Most experiments used a transgenic mouse line expressing Cre from the melanopsin (*Opn4)* locus of a BAC transgene (*Opn4^BAC-Cre^*; endogenous *Opn4* alleles were not targeted)^36^. Transduction of these mice with Cre-dependent viral vectors gave robust labeling of ipRGCs. However, crossing these mice with *ROSA26^FLEX-NL^* did not. Experiments on M4 ipRGCs used a line in which the gene encoding Cre recombinase replaced one allele of *Opn4*^20^. It was combined with *ROSA26^FLEX-NL^* (*Opn4^Cre/+^*; *ROSA26^FLEX-NL/+^*). Loss of one *Opn4* allele is inconsequential in these experiments because they concerned rod-driven thresholds, which this study found to resemble those of wild-type M4 ipRGCs^21,37^. Replacing both alleles of melanopsin (*Opn4^Cre/Cre^*) produced melanopsin-null mice. Experiments with rod or cone inactivation used *Gnat1^−/−^* or *Gnat2^−/−^* mice, respectively^38,39^. These knockouts disable phototransduction, locking rods and cones in the dark state without degeneration. All mouse lines were maintained on a C57BL/6J background. Mice of both sexes were used. Ages were P30-P180.

### Generation of nano-lantern viruses

To generate an AAV vector for Cre-dependent nano-lantern (NL) expression (AAV-FLEX-NL), the NL cDNA was PCR amplified from a pcDNA vector. It was subcloned into an AAV vector, inverted, between alternating loxP and lox2272 sites to generate a FLEX cassette. The construct was flanked by ITR sequences and contained an EF1α promoter, WPRE sequence, and hGH polyadenylation signal. To generate a Cre-independent AAV vector for NL expression (AAV-NL), loxP and lox2272 sites were removed and inversion of the NL cassette reversed (Genscript). Both vectors were validated with sequencing. Viruses (AAV2/1) were produced by the viral core of the Intellectual and Developmental Disabilities Research Center of Boston Children’s Hospital.

### Stereotaxic injection of viruses

Mice (>P60) were anesthetized with isoflurane, fixed in a stereotaxic frame, and prepared for sterile surgery. A midline incision revealed the skull and a craniotomy was made. *Opn4^BAC-Cre^* mice were injected with a 1:1 mixture of AAV2/1-FLEX-NL (4.4×10^13^ gc/ml) and AAV2/1-CAG-H2B-tdTomato (Addgene #116870, 2.4×10^13^ gc/ml); nuclear-localized tdTomato was used to check the injection site. In experiments using *Gnat1^−/−^* or *Gnat2*^−/−^ mice (which did not express Cre), AAV2/1-NL was used instead (2.9×10^13^ gc/ml). Viral injections were targeted to the SCN with the following stereotaxic coordinates, all measured from bregma: 0.88 mm lateral, 0.46 mm posterior, 5.90 mm travel distance. These injections were performed at a slight angle (6° from vertical, pointing toward the midline) to avoid tracking into the third ventricle. Injection pipettes were pulled from filamented borosilicate glass (1.0-mm outer diameter) and had a ∼60-µm tip diameter. Once the pipette was lowered to its target, it was kept in place for 5-10 minutes before infusing a total volume of 500 nl at a rate of 0.1 µl/min. After infusion, 5-10 min was given before slowly withdrawing the pipette from the brain. The wound was closed, mice recovered from surgery, and meloxicam or Ethiqa XR was given for analgesia. Viral expression was first visible ∼2 weeks following injection.

### Electrophysiology solutions

The extracellular solution was bicarbonate-buffered Ames’ medium (Sigma or US Biological) or “ionic” Ames’ medium^13,17^ (in mM: 120 NaCl, 22.6 NaHCO_3_, 3.1 KCl, 0.5 KH_2_PO_4_, 1.2 CaCl_2_, 1.2 MgSO_4_, and 6 glucose), both equilibrated continuously with carbogen (95% O_2_/5% CO_2_) for a pH of 7.4. Synaptic transmission was blocked with an established cocktail^11–13,16,17^: 3 mM kynurenic acid, 100 μM picrotoxin, 10 μM strychnine, and 100 μM D,L-AP4. The internal solution for perforated-patch recording was (in mM) 110 K-methanesulfonate, 13 NaCl, 2 MgCl_2_, 10 EGTA, 1 CaCl_2_, 10 HEPES, 0.1 Lucifer Yellow, and 0.125 amphotericin B; pH 7.2 with KOH. Amphotericin B was dissolved in DMSO to make 100× aliquots, and stored in the dark at -20 °C for up to two weeks. A liquid junction potential of +7 mV has been corrected. The pipette solution for loose-patch recording was HEPES-buffered Ames’ medium (in mM, 140 NaCl, 3.1 KCl, 0.5 KH_2_PO_4_, 1.2 CaCl_2_, 1.2 MgSO_4_, 6 glucose, 10 HEPES; pH 7.4 with NaOH).

Coelenterazine h stock was made at 200× in 100% ethanol under infrared light, and stored in darkness at -80 °C for up to six months.

### Retina preparation

Mice were dark adapted for >1 hour prior to experimentation, and subsequent procedures took place under infrared illumination using infrared-visible converters that were head- and microscope-mounted. Dim, indirect red light was used sparingly as needed. Mice were deeply anesthetized with Avertin, enucleated, and euthanized. The retina was isolated at room temperature in carbogenated Ames’ medium (see above) and vitreous humor removed. The retina was flattened with peripheral cuts and mounted on a glass coverslip coated with poly-L-lysine, RGC side up. Occasionally, to increase the number of recordings per animal while minimizing total light exposure during experimentation, the retina was cut in half, each piece was mounted on a separate coverslip, and one was incubated in carbogenated Ames’ medium at room temperature until use.

### Cell imaging and identification

Before electrophysiological recording, the mounted retina was transferred to a perfusion chamber containing carbogenated Ames’ medium. The medium was replaced with 1 ml of carbogenated Ames’ medium containing 20-50 μM coelenterazine h. The chamber was immediately moved to the electrophysiology rig and superfusion with carbogenated Ames’ medium was initiated. Detectable NL bioluminescence decayed over tens of minutes. During this period, as many NL-positive cells as possible were identified using an EM-CCD camera (Hamamatsu ImageEM) and the inner limiting membrane over their somata removed mechanically. Recordings were only initiated once the bioluminescence from these cells was no longer visible. After the conclusion of recording from a given retina, NL fluorescence was used to observe cell morphology and dendritic stratification. M1 cell identity was confirmed by the presence of dendritic stratification exclusively in the OFF sublamina (∼40 µm below the ganglion cell layer). M4 ipRGCs were identified by their bioluminescence (in the *Opn4^Cre/+^*; *ROSA26^FLEX-NL/+^* mouse line) and large somata^40^. Control M4 and other α-RGCs were targeted for recording by their soma size under infrared illumination; at the conclusion of the flash threshold determination, a 500-ms pulse of light was delivered to determine the α-RGC subtype (ON or OFF, sustained or transient)^41^.

### Electrophysiology

For loose-patch recording, pipettes (2-5 MΩ) were wrapped with parafilm and seal resistances were 10-30 MΩ. Recordings were conducted in voltage clamp with a holding potential of 0 mV. Recordings were discarded if the spontaneous spike rate or spike amplitude became unstable. Cells were also excluded from synaptic threshold measurements if their threshold firing showed evidence of melanopsin involvement (i.e., if firing continued to increase for 1-2 seconds following the flash, making fast synaptic responses impossible to separate; 4/36 cells). All loose-patch recordings were performed at 35 °C. For perforated-patch recording, pipettes (3.5-6 MΩ) were wrapped with parafilm to reduce capacitance. Seal resistances were >10 GΩ. Series resistances (typically <60 MΩ) were monitored but not compensated. Perforated-patch recordings were discarded if the initial seal resistance was <10 GΩ, access resistance was >100 MΩ, holding current at -80 mV was unstable, or any of these parameters changed abruptly. All perforated-patch recordings were performed at 23 °C for additional stability.

### Pupillometry

Animals were implanted with a headpost^42^, given a >1 week recovery period, and acclimated to the experimental setup. For experiments, they were dark adapted for >2 h and all subsequent procedures were done using dim red headlamps that were not directed toward the mouse. 1% tropicamide drops were applied to the stimulated eye to dilate its pupil, but not to the contralateral eye. Light was delivered by a Ganzfeld sphere (custom, internally coated with BaSO_4_), whose output is diffuse and thus allows precise quantification of retinal illumination^16^. The sphere was placed over the stimulated eye. An infrared light and video camera were focused on the contralateral eye to measure the consensual pupil response.

Optical stimuli matched those used *ex vivo* to measure duration-firing relations for 4.85 log R*/rod (at the retina) delivered in single pulses. Instead of 30 ms, 24 ms was used due to technical constraints, but both are instantaneous (impulse stimuli) for the retinal photoreceptors. The light intensity was scaled 7.14-fold to accommodate attenuation by the eye’s optics^16,43^. Stimuli were delivered in order of increasing intensity (and descending duration). Each stimulus duration (300 s, 30 s, 3 s, <30 ms) was delivered twice to each animal. The interstimulus interval was set to allow complete pupil dilation, assessed by the experimenter and then verified in analysis.

Recordings were analyzed in ImageJ using custom macros^16^. Recordings were adjusted for contrast and then binarized. The pupil was delineated by an ellipsoidal region of interest (ROI), whose maximum diameter was measured in each frame. Frames were inspected and those lacking accurate measurement were removed. Diameters were normalized to the mean in the 10-s before stimulus onset. Pupil diameters were averaged across trials and animals for each stimulus, binned (1 s), and used to calculate pupil area. This measurement was used to generate ‘pupil-corrected’ versions of the single-pulse temporal integration stimuli, in which the original stimulus intensity was multiplied by the normalized pupil area over time (**Fig. 6a-c**).

### Optical stimulation

Light sources were focused on the end of a liquid light guide for delivery to the microscope or the Ganzfeld sphere. For *ex vivo* experiments, light was collimated, sent through the microscope’s light path, focused by an objective or condenser, and centered on the soma of the recorded cell. For *in vivo* experiments, the light guide was positioned outside the animal’s field of view, such that all light entering the pupil had been internally reflected within the sphere.

All light sources were calibrated regularly using a radiometer. For broadband illumination, the spectrum’s shape was measured using a spectrometer and the intensity was scaled according to measurements with a radiometer. This procedure was used because the spectrometer was fiber-coupled and thus sensitive to the geometry of the stimulus, while the radiometer used a large, flat sensor that was insensitive. The scaling was performed by measuring the 460-nm band (10-nm width at half-maximum) with the radiometer and then scaling the spectrum accordingly, taking into consideration the attenuation within this band by the bandpass filter. Pulsed stimuli from LEDs were also verified with a fast photodiode.

#### Dim flashes

For eliciting threshold synaptic and melanopsin responses, light from a 75-W xenon arc lamp was filtered to select wavelength and intensity, excluding infrared and ultraviolet light. Stimulus timing, including for fine intensity adjustment (20-30 ms), was controlled by an electromechanical shutter. The beam was delivered through a 40× or a 4× objective to yield a spot diameter of 300 or 2800 µm, respectively.

#### Intensity-firing and duration-firing relations

For measuring intensity-firing relations, light from a 460-nm LED (Luminus PT120 or PT121) was bandpass filtered (460-nm center wavelength, 10-nm width at half-maximum). For duration-firing relations, a white LED was used (Luminus CBT140W). Both stimuli were collimated and delivered evenly over the whole retina using a substage condenser (as trans-illumination). Intensity was controlled with motorized neutral density filters (switching time <50 ms) as well as analog current and pulse-width modulation, using an Arduino Uno board and the ArduinoIO package for MATLAB.

### Optical stimulus quantification

Stimulus intensities are often reported in units of photoactivated rhodopsin molecules per rod (R*/rod) for flashes, and R*/rod/s otherwise. The conversion from photons to R* used an effective collecting area^37^ of 0.5 µm^2^ and a standard spectral template^44^ and rhodopsin’s λ_max_^45^ (502 nm). For a broadband stimulus, its spectrum was multiplied by the rhodopsin template, the product integrated, and the integral multiplied by the effective collecting area. For a narrowband stimulus (10 nm width), all photons were assumed to be of the center wavelength. The intensity was multiplied by rhodopsin’s relative sensitivity at this wavelength, then multiplied by the effective collecting area. If another photopigment was considered, its template was used in place of rhodopsin’s. The λ_max_ values used were 358, 508, and 471 nm for short-wavelength-sensitive cone photopigment, medium-wavelength-sensitive cone photopigment, and the ground (R) state of melanopsin^17,45–47^. For comparison with the behavioral experiments of Nelson and Takahashi^5^, their stimulus wavelength (503 nm) was referenced to the spectral template of rhodopsin (502 nm) and multiplied by the 0.5-µm^2^ effective collecting area.

Flashes and steps of light were compared using the integration time (t_i_) of the phototransduction cascade in question: 200 ms and 7.6 s for rhodopsin and melanopsin near body temperature, respectively, and 21.7 s for melanopsin at room temperature^16,17,48^. Flash intensities (photons/µm^2^) were divided by t_i_ to yield equivalent step intensities, and step intensities were multiplied by t_i_ to yield equivalent flash intensities^49^.

When measuring melanopsin dim-flash responses, monochromatic stimuli were reported in terms of log photons/µm^2^ rather than log R*/rod; this value is expressed at 471 nm, the λ_max_ of melanopsin’s ground state.

### Optical stimulus protocols

#### Synaptic flash thresholds

A 4× objective was used to ensure coverage of the entire dendritic arbor of M1s and the larger M4s/ON-sustained α-RGCs^40,50^. 410 nm was used because it activates all photopigments in the mouse retina similarly^17,45,46^. Intensities were presented in ascending order (4 repetitions at each intensity, with more given if a response was not immediately apparent). Trials were analyzed offline to find the minimum flash intensity that produced a detectable response for each cell within a 500-ms window.

#### Threshold spectral sensitivity

M1s were probed near threshold with flashes of varying intensity at 440 and 560 nm. These flashes were delivered through a 40× objective to allow corrections of cell and pipette positions, promoting recording stability, and because the spot suffices to cover the smaller M1 dendritic field^50^. Each trial consisted of 10-40 sweeps at the same flash intensity and wavelength, and wavelengths were alternated between trials. Flashes were calibrated to elicit the smallest reliable responses, usually with peaks of 2-5 pA. The average response amplitude for each trial was calculated, and the 440- and 560-nm flashes that produced the most similar responses were selected (they were within 15 ± 19%, which is 0.87 ± 1.2 pA; 20 cells). Typically, 1 or 2 matched response pairs were obtained for each cell. For each wavelength, dividing the response peak amplitude by the flash intensity gave sensitivity.

We expected a 440:560 sensitivity ratio of 1.34 for rhodopsin (peak wavelength sensitivity, λmax, of 502 nm) and 0.91 for medium-wavelength-sensitive (M) cone photopigment (508 nm; **Fig. 1b**); short-wavelength-sensitive (S) cone pigment (358 nm) absorbs these wavelengths poorly (10^−3^ and 10^−7^ of maximum, respectively) and should be negligible at the intensities tested^45,46^. Any melanopsin activation (471 nm for the ground state, the only one detectable after dark adaptation)^17,51^ should be clear, with a 440:560 nm ratio of ∼20 and distinctly prolonged kinetics^13,16,17,47^.

#### Measuring synaptic and intrinsic responses at threshold and above

The initial phase of these experiments was identical to those for threshold spectral sensitivity (see above), but 500-nm flashes were used. The lowest flash intensity that produced a clear response was identified, and responses obtained for averaging. The flash intensity was then increased by 1000× and the sweep length increased to 20 s to accommodate the slower responses that were often elicited. After measuring these responses, flashes of increasing intensity (500 or 480 nm) were given until peak melanopsin responses of 2-15 pA were visible; these responses were likely in the linear range and thus readily analyzed^13,16^. Occasionally, the synaptic responses near and above 1000× threshold were very small and resolved soon enough (within ∼2 s) that the peak of a near-threshold melanopsin response was distinguishable (e.g., cell E19021501B in **Fig. 1h**); in these cases, melanopsin’s dim-flash sensitivity was measured directly from this peak. Otherwise, synaptic antagonists were applied before probing flash intensities above 1000× threshold.

#### Intensity-firing relations

The first intensity staircase in the series consisted of 8 ascending steps, from -0.9 to 4.1 log R*/rod/s in ∼5-fold increments, of 30-s duration. The “steady” firing rate was measured in the last 10 s of each step. While longer steps (e.g., 120 s) allow M1s to reach a more consistent steady state^11^, 30-s steps allowed 2-3 staircases to be presented during a stable recording. Subsequent staircases were delivered with antagonists of synaptic transmission and after ≥10 minutes of dark adaptation, which sufficed for intrinsic responses to recover from the first staircase (**Extended Data Fig. 4**). These staircases were identical to the first but continued for 3 additional steps to reach up to 4.8-6.3 log R*/rod/s (spanning the intrinsic dynamic range of many M1s)^11^. These additional steps were excluded from the first staircase because they extended the duration of dark adaptation to beyond the stable lifetime of most recordings (35 °C). Of the 22 cells that were given the highest intensity step during the second staircase, 15 entered depolarization block, consistent with previous measurements^11^.

In synaptic antagonists, two cells did not reach the ≥1 Hz steady firing threshold within the tested stimulus range (see **Extended Data Table 3**, right column, and high-synaptic cluster in **Fig. 2b**). In these cases, thresholds were treated as right-censored data for statistical purposes (that is, as >I_max_, the maximum intensity delivered) and for visualization (see ΔI_θ_ values in **Fig. 2b**). Additional discussion is below, in **Statistics**.

#### Duration-firing relations

Stimuli for temporal integration experiments consisted either of single pulses (300 s, 30 s, 3 s, 30 ms) or of a set of distributed square pulses (0.1 Hz/0.1 duty cycle or 1 Hz/0.1 duty cycle, given as 30 or 300 pulses across 300 s). Each of these stimuli was calibrated to deliver the same total number of photons per unit area. The standard R* count was 4.85 log R*/rod (equivalent to 6 log photons/µm^2^ at the cornea)^16^. We chose this count for two main reasons. First, dividing it by the tested durations gives a range of intensities in which synaptic and intrinsic drives are likely to contribute. The shortest duration is roughly equivalent to moderate daylight (6.38 log R*/rod/s) and the longest to twilight (2.38 log R*/rod/s). Second, these intensities are subsaturating for circadian phase shifts. Also tested was 5.85 log R*/rod (7 log photons/µm^2^ at the cornea)^16^. Stimuli were delivered to each cell in random order, with intervening dark adaptation periods of ≥5 min. For each trial, the pre-stimulus firing rate was subtracted from the response rate.

### Immunohistochemistry

Phosphate-buffered saline (PBS) was the vehicle for all reagents. Retinas were fixed in 4% paraformaldehyde (30 min at room temperature), washed, blocked in 2.5-10% serum with 1% Triton X-100 (1 h at room temperature), and exposed to rabbit anti-GFP at 1:500 (Thermo Fisher A11122). Following incubation at 4 °C for 1-3 days, retinas were washed and then exposed to chicken anti-rabbit conjugated to Alexa 488 (Thermo Fisher A21441). A final wash was followed by mounting on slides with antifade medium containing DAPI. Retinas were imaged using a confocal microscope (Zeiss LSM510 or LSM710).

### Analysis

#### Statistics

Nonparametric statistical tests were used unless otherwise noted. Correlations were quantified as Spearman’s rank coefficients. To calculate statistical significance, the two-tailed Mann-Whitney U test was used for unpaired data and the two-tailed Wilcoxon signed-rank test was used for paired data. Paired data typically took the form of repeated measurements from the same cell within a recording; cells with missing values (due to recording lifetime) were not considered during significance testing. Significance testing involving right-censored data used a paired Prentice-Wilcoxon test, which is suited to censored values. Bonferroni corrections were applied when multiple comparisons were made among the same data sets; otherwise, p < 0.05 was considered significant. For k-means clustering (k = 2), Euclidean distance was used and classes were initialized using a random population subset. Z-score standardization of clustering variables made no difference in cluster assignments. When clustering variables included right-censored values (2/32 cells), the censoring limit was used.

#### Estimates of flash threshold for spiking

Flash intensities tested with <5 trials were omitted. For the remainder, spikes were counted in 100-ms windows, selected from the 500 ms before and after the flash, that contained the maximum and minimum spike counts. Baseline spike rates were subtracted. Three approaches were taken to estimate threshold. The first took the lowest flash intensity at which the pre- and post-flash spike counts appeared different (p = 0.05, one-tailed Wilcoxon signed-rank test, Pratt method for handling ties). The second used Receiver Operating Characteristic (ROC) analysis. For each flash intensity, the likelihood of observing *n* spikes after the flash was plotted against that of observing *n* spikes before the flash; varying n gave the ROC curve. The area under each curve was calculated and threshold taken as the lowest flash intensity at which this area was ≥0.60. The third approach was to represent each flash trial as a vector with a value of 1 at each spike time and 0 otherwise, then sum the vectors for every trial of a given flash intensity. This sum was convolved with a Gaussian with a standard deviation of 25 ms (the analysis was robust to reasonable variations of this parameter). Threshold was taken as the lowest flash intensity at which the peak of the convolution was ≥2 standard deviations above the mean, within 0.5 s of flash onset. Thresholds measured by these three methods were comparable. The last was used because it provided the closest match to visual inspection of the data.

#### Comparisons of duration-firing relations

Permutation tests were used to compare the population duration-firing (D-F) relation in single-pulse, temporal integration experiments. To compare two experimental groups (e.g., 4.85 log R*/rod with and without pupil correction; see **Fig. 6**), each cell’s D-F relation was randomly assigned to one of those two groups, keeping the group size consistent with the empirical data set. Single exponentials were then fit to the population D-F relation created from those shuffled data. Trials were excluded when the curve fit failed for either population D-F relation; that is, when either the fit’s Χ^2^ value exceeded 0.01, or when there was no change in Χ^2^ after 10 iterations. Each fit yielded a time constant, τ. The difference in τ between the two shuffled population D-F relations was compared to the difference observed empirically. 10,000 shuffles were used. The p value is the fraction of trials in which the difference in τ between shuffled population D-F relations exceeded the empirical value. These p values were 4.85 log R*/rod vs. pupil-corrected 4.85 log R*/rod, 0.0031; 4.85 log R*/rod vs. 4.85 log R*/rod in synaptic antagonists, 0.42; and 4.85 log R*/rod vs. 5.85 log R*/rod, 0.0073.

#### Refitting of behavioral phase shift data from Nelson and Takahashi

In the original study^5^, behavioral data (phase shift in minutes vs. stimulus intensity in photons/cm^2^/s) were fit to a modified Naka-Rushton equation:

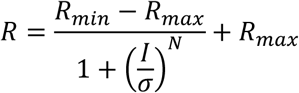

where *R* is the response, *R_min_* is the minimum response (0 by default), *R_max_* is the maximum response, *I* is the stimulus intensity, σ is the intensity of half-maximal response, and *N* is a cooperativity factor. To compare these data with cellular duration-firing relations, the raw, per-animal values were refit to this equation; published values of *R_min_* and *R_max_* were used, while *N* and σ were free parameters.

#### Software

Igor Pro (WaveMetrics) was used for curve fitting and most other analyses. Python was used in some cases, and R was used for the paired Prentice-Wilcoxon test.

